# The rates of introgression and barriers to genetic exchange between hybridizing species: sex chromosomes vs. autosomes

**DOI:** 10.1101/2020.04.12.038042

**Authors:** Christelle Fraïsse, Himani Sachdeva

## Abstract

Interspecific crossing experiments have shown that sex chromosomes play a major role in reproductive isolation between many pairs of species. However, their ability to act as reproductive barriers, which hamper interspecific genetic exchange, has rarely been evaluated quantitatively compared to Autosomes. This genome-wide limitation of gene flow is essential for understanding the complete separation of species, and thus speciation. Here, we develop a mainland-island model of secondary contact between hybridizing species of an XY (or ZW) sexual system. We obtain theoretical predictions for the frequency of introgressed alleles, and the strength of the barrier to neutral gene flow for the two types of chromosomes carrying multiple interspecific barrier loci. Theoretical predictions are obtained for scenarios where introgressed alleles are rare. We show that the same analytical expressions apply for sex chromosomes and autosomes, but with different sex-averaged effective parameters. The specific features of sex chromosomes (hemizygosity and absence of recombination in the heterogametic sex) lead to reduced levels of introgression on the X (or Z) compared to autosomes. This effect can be enhanced by certain types of sex-biased forces, but it remains overall small (except when alleles causing incompatibilities are recessive). We discuss these predictions in the light of empirical data comprising model-based tests of introgression and cline surveys in various biological systems.

## INTRODUCTION

Speciation is a process of gradual accumulation of reproductive barriers in the genome, ultimately leading to a cessation of gene flow between groups of individuals forming distinct biological species (Dobzhansky 1937). The extent to which barrier loci reduce interspecific gene flow is a central factor in the study of speciation, as it allows the strength of reproductive isolation to be quantified as populations diverge (see Ravinet et al. 2017 for a review). Another major advance in the genetics of speciation has been the discovery, by interspecific crossing experiments, of two extremely robust patterns: (i) “Haldane’s rule” (Haldane 1922; Schilthuizen et al. 2011), i.e. in species with sex-specific reduced fitness of Fl hybrids, the affected sex is generally heterogametic; and (ii) the “large-X/Z effect” (Dobzhansky 1936), i.e. the disproportionate density on the X or the Z chromosome of barrier loci causing hybrid sterility or inviability. These “rules of speciation” (Coyne and Orr 1989) suggest the existence of universal mechanisms associated with sex chromosomes that may promote speciation. However, these two lines of research (quantifying the barrier strength vs characterizing the genetic basis of speciation) have, for the most part, been conducted independently. In particular, there is room for improvement in the empirical estimation of the barrier to interspecific gene flow due to specific chromosomes, as this information is mainly available only for model species (Payseur et al. 2018). In consequence, the ability of sex chromosomes to act as barriers that impede interspecies gene exchange has been little evaluated quantitatively or in a systematic way.

The recent explosion of genomic data in the field of speciation has provided empirical evidence across a diversity of species with different sex chromosome systems (i.e. XY under male heterogamety, and ZW under female heterogamety, Bachtrog et al. 2014). Indeed, the presence of reproductive barriers along the genome reduces interspecific gene flow around each barrier locus, leading to a local increase of interspecific differentiation relative to the rest of the genome. This effect has been widely used to scan genomes for regions that are abnormally differentiated between species, in search of loci involved in speciation. These studies collectively show a systematically stronger differentiation of sex chromosomes (X or Z) compared to the autosomes (Presgraves 2018), even after correcting for the different effective population sizes of sex chromosomes and autosomes. This indicates that sex chromosomes may play a leading role in the isolation of nascent species. However, caution must be exercised when interpreting this pattern, as measures of relative differentiation (such as *F*_*ST*_; Weir and Cockerham 1984) are sensitive to the level of diversity within species, and whether or not gene flow occurred during species divergence (Charlesworth 1998; Cruickshank and Hahn 2014). Complementary approaches addressing this limitation are based on simple summary statistics of absolute divergence (such as *D*_*XY*_; Nei and Li 1979), cline analyses (Barton and Hewitt 1985) and speciation model inferences (Sousa and Hey 2013), but they do not reveal clear-cut patterns (see Discussion and Table 3).

Three main theories have been proposed to explain why sex chromosomes might act as stronger interspecies barriers than autosomes. First, sex chromosomes may accumulate interspecific barrier loci faster than autosomes. The faster-X theory (Charlesworth et al. 1987) predicts that this would be the case if incompatible loci first appear as recessive or partially recessive beneficial mutations (or under a much wider range of dominance levels, if the effective population size for the X relative to autosomes is high enough; Vicoso and Charlesworth 2009). This is because the hemizygosity of the sex chromosome, i.e. the fact that the heterogametic sex has only one copy of the X/Z, makes selection more efficient by unmasking sex-linked recessive mutations. Most theoretical and empirical work so far deals with the faster-X theory; however, empirical evidence in favor of greater efficacy of selection on the sex chromosome is mixed (Meisel and Connallon 2013; but see Charlesworth et al. 2018, who provide fairly solid evidence for adaptive faster-X effects in *Drosophila).* Instead, other studies have suggested that the accumulation of slightly deleterious mutations may be a major cause of faster-X/Z, especially in birds, where the variance of male reproductive success strongly decreases the effective population size of the sex chromosome relative to autosomes, increasing genetic drift (Mank et al. 2010).

Another theory, which was initially dismissed but has recently returned to the limelight, is that barrier loci may occur more readily on the sex chromosomes, because the absence of recombination between the X and the Y (or the Z and the W) makes them more susceptible to segregation conflicts in the heterogametic sex (e.g. Bachtrog et al. 2019). Since segregation distorters and their suppressors co-evolve within populations independently, changes in these interactions in hybrids may cause incompatibilities (Frank 1991; Hurst and Pomiankowski 1991; and see Phadnis and Orr 2009, for an example of a X-linked meiotic driver associated with hybrid sterility in *Drosophila).* This mechanism thus contrasts with the classical view that interspecies incompatibilities first appear within species as beneficial mutations (Presgraves 2010).

The last theory does not require that sex chromosomes harbor more barrier loci than autosomes, but simply that they are more exposed to selection, such that sex-linked incompatibilities cause more deleterious effects (than autosomal incompatibilities) in hybrids. This process has been formalized by the dominance theory (Turelli and Orr 1995, 2000), which is also based on the hemizygosity of the sex chromosome and assumes that most incompatible mutations act recessively in hybrids. However, it has been experimentally tested (and validated) almost exclusively in *Drosophila* (e.g. Masly and Presgraves 2007; Cattani and Presgraves 2012; Llopart et al. 2018, but see Matsubara et al. 2015 for a study in rice). Assessing its generality remains difficult, due to the experimental challenge of measuring the dominance of mutations.

The present work aims to understand the role of the X or Z chromosome (note that we do not consider the Y or W) in the formation of new biological species by focusing on the impact of sex-linked reproductive barriers on the reduction of gene flow between hybridizing species, rather than on their rate of establishment during species divergence. To do this, we extend the theoretical framework of Barton and Bengtsson who modeled the introgression of incompatible autosomal genomic blocks between hybridizing species (Barton 1983), and quantified their effect on the effective migration rate at a linked neutral marker (Barton and Bengtsson 1986). With multiple barrier loci, Barton and Bengtsson showed that neutral gene flow is significantly reduced over most of the genome only if the number of loci involved in reproductive isolation is large enough that any neutral marker becomes closely linked to at least one barrier locus. While they showed that this result is little influenced by the position of the neutral marker along a given chromosome, the effect of being located on the sex chromosomes rather than on autosomes was not addressed.

Yet, for the above reasons, one may expect to see differences between the two types of chromosomes: in the heterogametic sex, the greater efficacy of selection against deleterious mutations and the lack of recombination over much of the X or Z are expected to reduce the rate of introgression of sex-linked neutral markers relative to autosomal ones, even when there is no difference between the number, density or selective effects of sex-linked and autosomal incompatibilities. Moreover, because sex chromosomes are involved in sex determination, they are inherited differently between males and females and thus spend different periods of time in each sex, making them sensitive to sex-specific evolutionary forces (Hedrick 2007), such as sex-biased migration, sex-biased selection, or achiasmy (the absence of recombination in the entire genome of the heterogametic sex, which causes the X/Z to have more recombination than the autosomes; e.g. see Betancourt et al. 2004). Since these evolutionary forces determine the intensity of the barrier to gene flow (Barton and Bengtsson 1986), their sex specificity must be taken into account in order to quantify the role of sex chromosomes in speciation. An additional complication comes from the phenomenon of dosage compensation that has been found in many species with a degenerated sex chromosome (Mank 2013). The degeneration of the Y (or the W) causes a dose difference between the sexes, so that sex-linked loci are transcribed half as much in the heterogametic sex. Dosage compensation then consists of restoring similar levels of the X (or Z) gene product in males and females. Among the diversity of mechanisms that have evolved (Gu and Walters 2017), some may lead to stronger selection against hybrids, and thus should enhance the ability of sex chromosomes to act as interspecific barriers.

To date, only a handful of theoretical studies have examined the role of sex chromosomes in speciation. The dominance theory (Turelli and Orr 1995, 2000) offered a simple explanation for the two “rules of speciation”, building upon the allopatric model of postzygotic isolation elaborated by Dobzhansky (1937) and Muller (1940). This seminal model involves deleterious epistasis between alleles at different loci (i.e., Dobzhansky-Muller incompatibilities, or DMIs). In a model of parapatry, Hoellinger and Hermisson (2017) found that X-linked DMIs are more easily maintained in the presence of interspecies gene flow than autosomal DMIs. All these studies address the question of the origin, maintenance or accumulation of incompatibilities, but do not explicitly quantify their long-term effect as barriers to neutral gene flow. In fact, it may be that DMIs are generally ineffective at maintaining genome-wide differentiation in the face of gene flow (while they can create strong selection against Fl and F2 hybrids) as shown by simulations (Lindtke and Buerkle 2015).

In contrast, Muirhead and Presgraves (2016) modeled the effect of a locally deleterious mutation (either intrinsically incompatible with the local genome, or extrinsically incompatible with the local habitat) on gene flow at a linked neutral marker. Under a weak migration/strong selection regime, they showed that selection against incompatible mutations reduces the flow of neutral markers more strongly on the sex chromosomes than on autosomes, the effect being greater when mutations are recessive.

While all these studies are in line with a reduced introgression probability on the sex chromosomes relative to autosomes, they are limited to single/pairs of barrier loci, even though speciation is a complex, and probably multigenic, process. Thus, it is essential to develop more realistic models that account for the effects of multiple linked barrier loci. This can have important implications; for example, Barton (1983) showed that linkage between incompatible autosomal alleles, which governs the extent to which they act as a single unit of selection, is a key determinant of their level of introgression and the barrier to gene flow they induce. A multilocus theory would thus allow us to better understand the differences between sex-linked and autosomal interspecies barriers, and the causes underlying these differences, such as the role of recombination. In addition, these models provide predictions of introgression patterns along chromosomes, and thus are critical to capture the signature of polygenic barriers in genomic data.

Here, we address this gap by extending previous multilocus predictions (Barton 1983; Barton and Bengtsson 1986) to the case of sex chromosomes. We introduce a model where chromosomes contain multiple interspecies incompatibilities, and quantify (i) their frequency on sex chromosomes relative to autosomes in populations under migration-selection balance and (ii) the effect of selection against these incompatibilities on the neutral gene flow between two hybridizing species. Our aim is to understand (i) to what extent the particularities of the sex chromosomes: hemizygosity and lack of recombination on the X/Z in the heterogametic sex (we ignore pseudoautosomal regions) affect their ability to hinder interspecific gene flow, and (ii) the influence of sex-biased processes and dosage compensation in different sexual systems (XY and ZW). Our model is general in the sense that it applies to both male heterogamety (males are XY and females are XX) and female heterogamety (females are ZW and males are ZZ), and it allows us to address the effects of various kinds of sex-specificity by introducing sex-specific parameters. Unless otherwise stated, we assume an XY system, as the results apply to ZW systems by interchanging males and females.

## MODEL AND METHODS

We consider a population with males and females. The homogametic sex carries two copies of the sex chromosome (X or Z) and two copies of each autosome; the heterogametic sex carries one copy of the sex chromosome and two copies of each autosome. We focus on X/Z chromosomes (or regions of them) where there are no homologous genes on the Y/W, and which do not undergo recombination in the heterogametic sex.

Hybridisation between two incompletely isolated species occurs in a mainland-island setting. Migration is assumed to be one-way from a donor to a recipient species, although our results hold for two-way exchange as long as migration is weak and introgressed alleles rare. We model the introgression of a single medium-sized block of genome with multiple loci that influence fitness in the recipient species. We assume that the two species are fixed for different alleles at these loci, and that each introduced allele is deleterious in the recipient species, either because it is not adapted to the local environment or to the local genetic background. Throughout we neglect *de novo* mutations and segregating variants (with effects on fitness) that may be private to one of the two species. We also neglect the effect of loosely linked or unlinked incompatible alleles (spread across the entire genome), since these would affect introgression of both autosomal and X-linked blocks to a similar extent via an increase in the fitness variation and a resultant reduction in the effective population size (Barton 1995). We thus focus here on a comparison of the introgression dynamics of a single X-linked vs autosomal block.

In each generation, a fraction *m*_*F*_ of females and a fraction *m*_*M*_ of males are replaced by immigrant females and males respectively. Immigrants are drawn from a donor species which is fixed for the deleterious block. Thus, if the block is on the X, immigrant females introduce two identical copies and immigrant males a single copy into the recipient species, while if it is autosomal, any immigrant introduces two copies of the deleterious block. The block is assumed to carry *L* equally spaced loci with deleterious alleles of equal effect, which multiplicatively reduce individual fitness in the recipient species. Thus the relative fitness of a female heterozygous for *y* and homozygous for *y′* X-linked introgressed alleles is 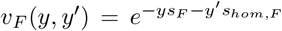, while the fitness of a male who carries *y* introgressed alleles is 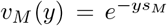. Here, *s*_*F*_ and *s*_*hom,F*_ denote heterozygous and homozygous effects per allele in females, and *s*_*M*_ the hemizygous effect in males; unlike in the more standard notation where *hs* refers to the heterozygous effect. Our choice of notation is motivated by the fact that, to the first order in *sL*, introgression depends only on heterozygous effects and is independent of the dominance coefficient, *h* (see below). Fitness in the autosomal case can be defined analogously. The per generation rate of recombination between adjacent selected loci is denoted by *c*_*F*_ in females (for either autosomal or X-linked blocks) and *c*_*M*_ in males (only for autosomes; no recombination occurs between the X and Y in males). Unless stated otherwise, we always assume autosomal recombination in the heterogametic sex.

We also analyze the introgression of a neutral marker linked to the deleterious genomic block. For simplicity, we focus on the so-called “rod” model of Barton and Bengtsson (1986), where the neutral marker lies at one end of the block, with the rate of recombination between the neutral marker and the terminal selected locus *αc*_*F*_ in females and *αc*_*M*_ in males. Here *α* parameterizes the map distance between the neutral marker and nearest selected locus, relative to the interlocus map distance. Thus, in this model, the neutral marker can escape into the recipient background via a single recombination event. Our analysis can be generalized to other marker configurations, following Barton and Bengtsson (1986) (e.g. the “embedded” model, Figure S7).

We derive analytical results by assuming that: (i) the frequency of the introgressing block (and its descendant fragments) is low so that matings between individuals carrying introgressed material can be neglected, and (ii) the deleterious block is short enough that multiple crossovers within the deleterious block can also be neglected. This implies that any individual male or female carries at most a single deleterious block. In particular, immigrant females (who carry two copies of the selected blocks) must necessarily pass on only one of these to their offspring (who are likely to inherit the other chromosome from a father bearing no introgressed material). Thus, in this regime, introgression is largely independent of the homozygous effect of deleterious alleles and is governed by the heterozygous (in females) and hemizygous (in males) selective effects.

In the following, we first present dynamical equations for the evolution of deleterious fragments of the introduced genome and solve these at equilibrium (i.e., migration-selection-recombination balance) to obtain analytical predictions for the equilibrium frequencies of deleterious X-linked blocks. We then calculate the effective migration rate of neutral markers into the recipient species, thus quantifying the reduction in gene flow at a neutral marker due to linkage to interspecific barrier loci. We also test these analytical expressions (i.e., the exact formulae and the weak-selection approximations) against individual-based simulations. We first analyze a basic model assuming co-dominance, multiplicative effects and no sex-specific forces, and then extend to sex-specificities, dominance and epistasis. Key notation is summarized in Table 1.

**Table 1.**
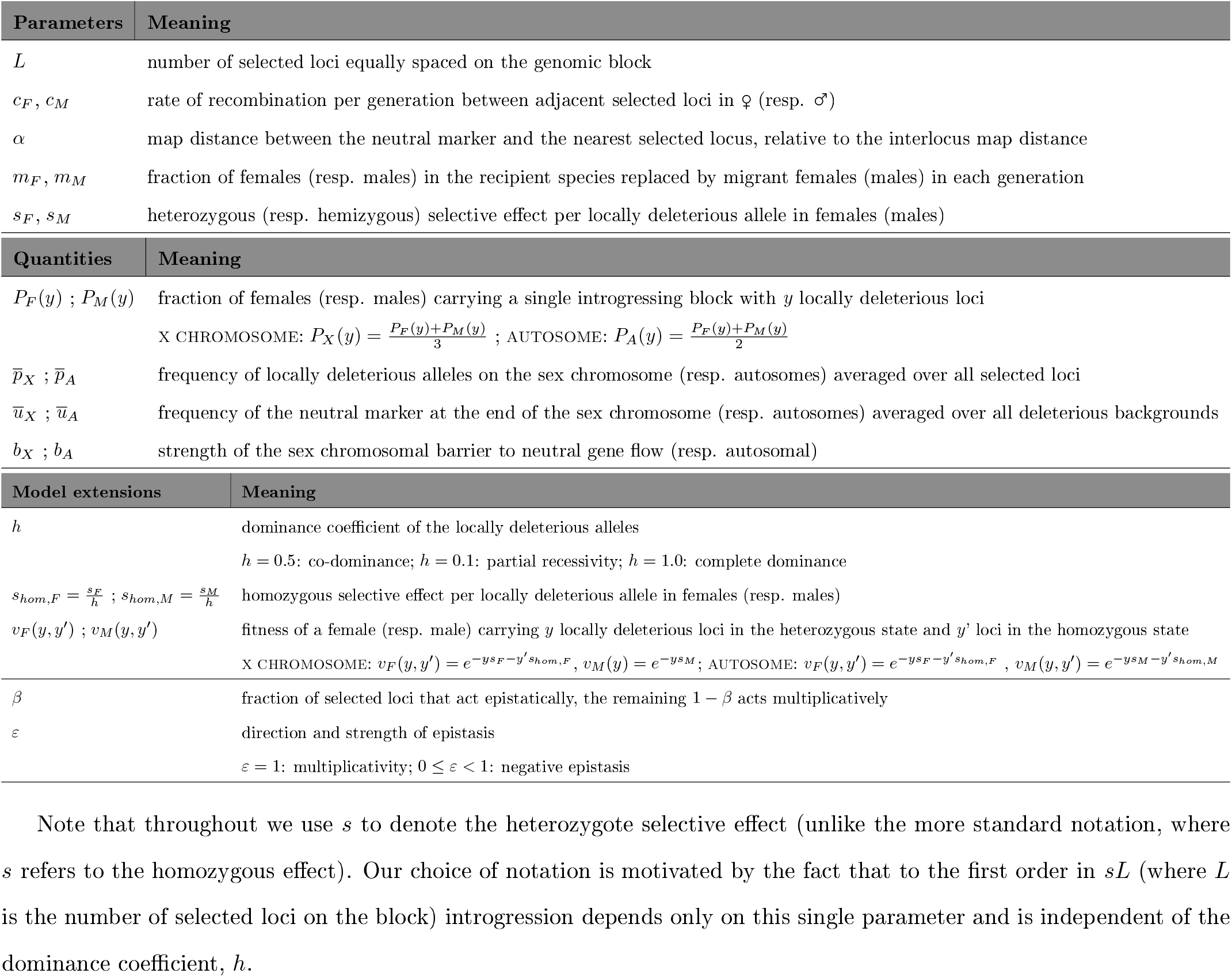
Notation.

| Parameters | Meaning |
| --- | --- |
| $L$ | number of selected loci equally spaced on the genomic block |
| $c_F, c_M$ | rate of recombination per generation between adjacent selected loci in ♀ (resp. ♂) |
| $\alpha$ | map distance between the neutral marker and the nearest selected locus, relative to the interlocus map distance |
| $m_F, m_M$ | fraction of females (resp. males) in the recipient species replaced by migrant females (males) in each generation |
| $s_F, s_M$ | heterozygous (resp. hemizygous) selective effect per locally deleterious allele in females (males) |
| Quantities | Meaning |
| $P_F(y) ; P_M(y)$ | fraction of females (resp. males) carrying a single introgressing block with $y$ locally deleterious loci |
| | X CHROMOSOME: $P_X(y) = \frac{P_F(y)+P_M(y)}{3}$ ; AUTOSOME: $P_A(y) = \frac{P_F(y)+P_M(y)}{2}$ |
| $\bar{p}_X ; \bar{p}_A$ | frequency of locally deleterious alleles on the sex chromosome (resp. autosomes) averaged over all selected loci |
| $\bar{u}_X ; \bar{u}_A$ | frequency of the neutral marker at the end of the sex chromosome (resp. autosomes) averaged over all deleterious backgrounds |
| $b_X ; b_A$ | strength of the sex chromosomal barrier to neutral gene flow (resp. autosomal) |
| Model extensions | Meaning |
| $h$ | dominance coefficient of the locally deleterious alleles |
| | $h = 0.5$: co-dominance; $h = 0.1$: partial recessivity; $h = 1.0$: complete dominance |
| $s_{hom,F} = \frac{s_F}{h} ; s_{hom,M} = \frac{s_M}{h}$ | homozygous selective effect per locally deleterious allele in females (resp. males) |
| $v_F(y, y') ; v_M(y, y')$ | fitness of a female (resp. male) carrying $y$ locally deleterious loci in the heterozygous state and $y'$ loci in the homozygous state |
| | X CHROMOSOME: $v_F(y, y') = e^{-y s_F - y' s_{hom,F}}$, $v_M(y) = e^{-y s_M}$; AUTOSOME: $v_F(y, y') = e^{-y s_F - y' s_{hom,F}}$, $v_M(y, y') = e^{-y s_M - y' s_{hom,M}}$ |
| $\beta$ | fraction of selected loci that act epistatically, the remaining $1 - \beta$ acts multiplicatively |
| $\varepsilon$ | direction and strength of epistasis |
| | $\varepsilon = 1$: multiplicativity; $0 \leq \varepsilon < 1$: negative epistasis |

Note that throughout we use $s$ to denote the heterozygote selective effect (unlike the more standard notation, where $s$ refers to the homozygous effect). Our choice of notation is motivated by the fact that to the first order in $sL$ (where $L$ is the number of selected loci on the block) introgression depends only on this single parameter and is independent of the dominance coefficient, $h$ .

### Introgression of multilocus deleterious blocks

#### Autosomal introgression

We consider how selection against multiple deleterious alleles influences introgression of an incompatible autosomal block in a regime where the vast majority of individuals carry at most one fragment of the block. While Barton (1983) considered the simplest case of haploid introgression and no sex differences, we generalize to the case where selection coefficients, migration rates and recombination rates may differ between males and females (see Appendix A, Supp. Text). For small parameters (i.e. *sL, cL, m* ≪ 1), we can take a continuous time limit by approximating *e*^−*sy*^ ≈ 1 − *sy* and, neglecting all terms that are second order in small parameters, e.g., *O*(*s*^2^), *O*(*ms*) etc. Then the frequencies *P*_*A*_(*y, t*) of introgressed fragments with *y* deleterious alleles at time *t* satisfy dynamical equations which are identical to those in Barton (1983) but with *s*_*A*_, *m*_*A*_ and *c*_*A*_ given by the sex-averaged rates 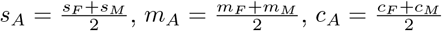; see Table 2):

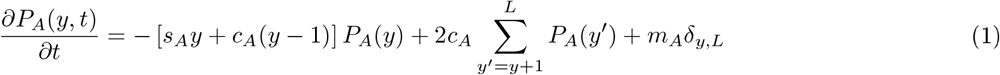

**Table 2.** Sex-averaged effective parameters.

| Parameters | Meaning |
| --- | --- |
| $s_X ; s_A$ | effective heterozygous selection per locally deleterious allele on the X chromosome (resp. autosomes) |
| | X CHROMOSOME: $s_X = \frac{2s_F+s_M}{3}$ ; AUTOSOME: $s_A = \frac{s_F+s_M}{2}$ |
| $m_X ; m_A$ | effective migration on the X chromosome (resp. autosomes) |
| | X CHROMOSOME: $m_X = \frac{2m_F+m_M}{3}$ ; AUTOSOME: $m_A = \frac{m_F+m_M}{2}$ |
| $c_X ; c_A$ | effective recombination on the X chromosome (resp. autosomes) |
| | X CHROMOSOME: $c_X = \frac{2c_F}{3}$ ; AUTOSOME: $c_A = \frac{c_F+c_M}{2}$ |
| $\theta_X ; \theta_A$ | strength of coupling between locally deleterious alleles on the sex chromosome (resp. autosomes) |
| | X CHROMOSOME: $\theta_X = \frac{s_X}{c_X}$ ; AUTOSOME: $\theta_A = \frac{s_A}{c_A}$ |
| $\frac{m_X}{s_X} ; \frac{m_A}{s_A}$ | expected allele frequency when selected loci are perfectly linked on the sex chromosome (resp. autosomes) |

These effective parameters govern introgression assuming that $sL$ , $cL$ and $m$ are $\ll 1$ .

Here *δ*_*y,L*_ equals 1 for *y* = *L*, and is zero otherwise. The first term represents the loss of the introgressed fragment due to selection against deleterious alleles and backcrossing with the recipient species; the second term corresponds to the creation of the fragment by splitting of larger fragments; the third term represents the influx of the full block (with *L* deleterious alleles) due to migration from the donor species. Note that eq. (1) is linear in *P*_*A*_*(y)* since we assume introgressing blocks to be rare and terms of the form *P*_*A*_(*y*)*P*_*A*_*(y′)* (which arise from mating between individuals carrying introgressing blocks) to be negligible. For the same reason, we can ignore terms of order 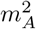; thus our results also hold for weak bidirectional migration.

Eq. (1) can be solved to obtain the block frequencies *P*_*A*_*(y)* at equilibrium and the average equilibrium frequency of deleterious alleles (averaged over all loci on the autosomal block): 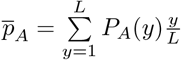 (see eq. (12) in Appendix A).

A key parameter governing introgression is 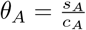, the ratio of selective effect to recombination rate per locus, which determines the extent of coupling between loci (Barton 1983). For large *L*, the average deleterious allele frequency 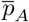 can be approximated as:

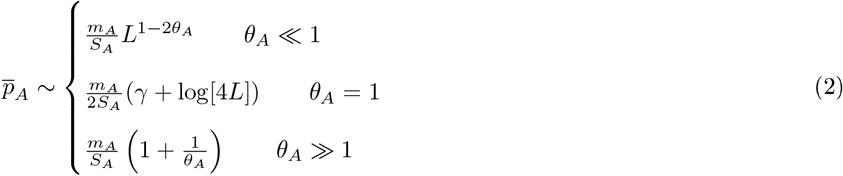

where *S*_*A*_ = *s*_*A*_*L* and *γ* = 0.577216 is Euler’s constant. One can define the effective selection coefficient, 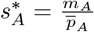 as the selective pressure at a single locus that would be needed to produce the observed average frequency of the deleterious alleles (Barton 1983). By extension, one can also define the effective number of loci 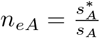, that, if fully linked, would have produced the same effect on deleterious allele frequency as the full deleterious block (Figure SI). Note that in the limit c → 0, i.e., for perfect linkage between deleterious alleles, *n*_*eA*_ must approach *L*.

For *θ*_*A*_ ≫ 1, (i.e., very strong coupling between loci), the deleterious allele frequency depends only on the net selective disadvantage *S*_*A*_ = *s*_*A*_*L* of the block, and is independent of the number of loci on the block (for large *L*). It also follows that the effective selection coefficient per locus, given by 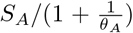, approaches the net selective effect *S*_*A*_ for very large values of *θ*_*A*_. In contrast, for *θ*_*A*_ ≪ 1 (i.e., weak coupling between loci), each locus experiences an effective selective disadvantage (given by 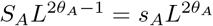) which is proportional to its own selective effect *s*_*A*_, but also depends weakly on *L* (the number of selected loci that are loosely linked to it). Crucially, in this regime, the effective selection per locus decreases and the deleterious allele frequency increases as we consider introgression scenarios where the same total selective effect is due to a larger and larger number of loci of weaker effect (i.e, on increasing *L* while keeping *S*_*A*_ = *s*_*A*_*L* and *C*_*A*_ = *c*_*A*_*L* constant). Thus for weak coupling between loci, this analysis (which assumes that deleterious allele frequency is low) remains valid only for small values of *L*, more specifically for 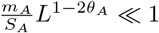.

#### X-linked introgression

We now analyze introgression of an X-linked block. Let *P*_*F*_*(y)* and *P*_*M*_*(y)* denote the fraction of females and males who carry a single copy of, i.e., are respectively heterozygous and hemizygous for, an X-linked introgressed block with *y* deleterious alleles. Thus, in a population of size *N* with a 1 : 1 sex ratio, the number of X-linked blocks with *y* deleterious alleles must be 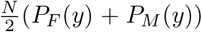, while the total number of X chromosomes is 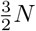. Thus, the frequency of X-linked introgressed blocks with *y* deleterious alleles is given by 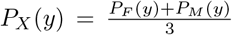. In Appendix A, we derive dynamical equations for the evolution of *P*_*F*_*(y,t)* and *P*_*M*_*(y, t)*, In the continuous time limit, i.e., assuming *sL*, *cL* ≪ 1, and *m* ≪ s, these can be combined into a single equation for the evolution of *P*_*X*_*(y, t)*:

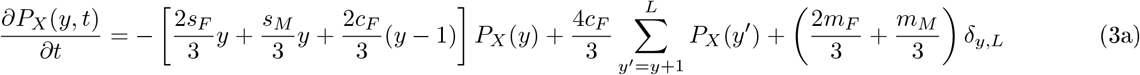

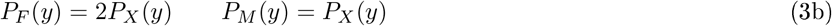

This equation is identical in form to eq. (1) (which describes the dynamics of the autosomal block distribution), but with *s*_*A*_ replaced by 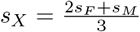, *m*_*A*_ by 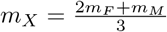 and *c*_*A*_ by 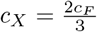 (see Table 2). The sex-averaged effective parameters *m*_*X*_, *s*_*X*_ and *c*_*X*_ are weighted sums of the corresponding male and female parameters. Female migration contributes twice as much as male migration to *m*_*X*_ simply because any female migrant introduces two X-linked blocks while a male migrant carries one. Similarly, the sex-averaged selection coefficient *s*_*X*_ is the sum of male and female components, with the contribution of *s*_*F*_, the heterozygous selective effect per allele in females, being twice that of *s*_*M*_, the hemizygous effect in males. Note that this is despite the fact that almost all females, with the exception of immigrant females, carry only one copy of any X-linked deleterious block (under the assumption of low migration). The 2:1 contributions emerge nevertheless because there are twice as many females who are heterozygous (for any X-linked deleterious block) as males who are hemizygous (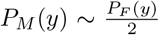, see eq. (3b)): this, in turn, is simply because females can inherit the block from either parent, while males only inherit it from their mothers. Thus any X-linked deleterious allele spends twice as much time in the heterozygous state in females than in the hemizygous state in males, causing it to be more sensitive to selection in females. The 2:1 contributions of female and male recombination rates (where the latter is assumed to be zero) to the sex-averaged recombination fraction *c*_*X*_ can be explained similarly.

Equation (3a) can be solved to obtain an explicit expression for the equilibrium *P*_*X*_*(y)* (details in Appendix A):

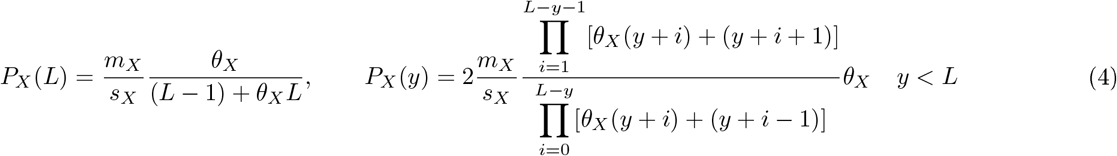

Here 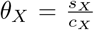 is a measure of the strength of coupling between deleterious alleles on the X chromosome. As in the autosomal case, equation (4) can be used to calculate the average frequency of deleterious alleles, averaged over all loci on the X chromosome: 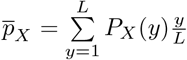. Since eq. (1) and (3) have the same form, this is given by the same expressions as in eq. (2), but with *s*_*X*_ and *m*_*X*_ replaced by *s*_*A*_ and *m*_*A*_ respectively. As before, we expect our analysis to become inaccurate when the X chromosome contains a large number of weakly linked and weakly selected deleterious alleles, i.e., for *θ*_*X*_ ≲ 1 and 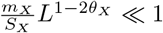.

Equations (1) and (3) reveal a simple correspondence between introgression on autosomes and on X chromosomes, even when there is multilocus selection against introgression (for weak selection and recombination: *sL,cL* ≪ 1). More specifically, they suggest that the distribution of lengths of X-linked introgressing blocks should be nearly identical to the distribution of autosomal introgressing blocks, upon comparing autosomes and sex chromosomes with 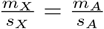 and *θ*_*X*_ = *θ*_*A*_ (this prediction is verified in simulations, see Figure S2A). Conversely, if we compare autosomal and X-linked blocks with identical genetic architectures and no sex-bias, then we have 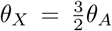 i.e., deleterious alleles on the X are more strongly coupled to each other (than on autosomes), resulting in lower rates of introgression. We quantify this precisely in the Results using our analytical expressions for *P*_*X*_*(y)* and *P*_*A*_*(y)* (eq. (4)).

Note that this correspondence between X-linked and autosomal introgression also applies to the dynamics of block length distributions (eq. (3a), see also Figure S2B for verification in simulations). This implies that the time-dependent distribution of block lengths can be described by the same mathematical results as for the autosomal case (Appendix in Baird 1995). Thus we do not consider dynamics any further here, and focus only on patterns at equilibrium.

### Multilocus barrier to neutral introgression

#### Autosomal barrier

We next investigate how deleterious genomic blocks act as genetic barriers to neutral gene flow by calculating the rate at which an allele at a linked neutral marker (fixed in the donor species) increases in the recipient species, where it is initially absent. We again assume that introgression is rare and that multiple crossovers within the deleterious block can be neglected. Therefore, the neutral marker can only be found in the recipient species associated with a consecutive series of between 0 and *L* locally deleterious alleles, and paired with a chromosome of the recipient species.

Following Bengtsson (1985), the strength of the barrier (*b*_*A*_) is defined as 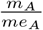, where *m*_*A*_ is the raw migration rate on autosomes and *me*_*A*_ is the effective migration rate defined as 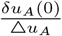. Here, Δ*u*_*A*_ is the difference in allele frequency of the neutral marker between the donor and recipient species (and is 1, by definition), while *δu*_*A*_(0) is the rate of increase of the neutral marker in the recipient background. The rate of increase *δu*_*A*_(0) can be calculated by tracking the rate at which the neutral marker is transferred between different genetic backgrounds (with different numbers of deleterious alleles) at equilibrium (Barton and Bengtsson 1986; see also Appendix B, Supp. Text). For *sL, cL* ≪ 1, the autosomal barrier strength *b*_*A*_ under this “rod” configuration can be written as (Barton and Bengtsson 1986):

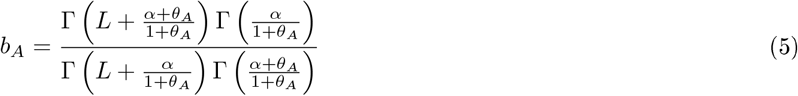

where Γ refers to the Gamma function.

The barrier strength is thus determined by the coupling coefficient 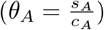 and the proximity of the neutral marker to the selected block (*α*). The dependence of the barrier strength on *θ*_*A*_ is qualitatively similar to the dependence of the equilibrium frequency of locally deleterious alleles. When *θ*_*A*_ is large (*sL* ≫ *cL*), all that matters in order for the neutral marker to introgress is to recombine away from the entire deleterious block in a single step, while when *θ*_*A*_ is small (*sL* ≪ *cL*), pairwise interactions between the neutral marker and each selected locus modulate neutral introgression.

Concerning the proximity of the neutral marker to the incompatible block, the smaller *α*, the stronger the barrier, since the neutral marker is then closely linked to the block. When *α* ≪ 1, the barrier strength is proportional to the effect of the closest deleterious locus, but the additional effect of the other deleterious loci on the block reduces gene flow well below the single-locus expectation. When *α* ≫ *L*, the deleterious block acts as a single unit with regard to the neutral marker, and thus the barrier strength is close to that produced by a single locus with total effect *sL* at a distance *ac* from the marker.

#### X-linked barrier

We now consider the case of a neutral marker on the X chromosome. We obtain an explicit expression for the strength of the barrier to neutral X-linked gene flow, under the same assumptions as in the autosomal case (see Appendix B for further details, Supp. Text). This is the same expression as in eq. (5), but with *θ*_*A*_ substituted by *θ*_*X*_:

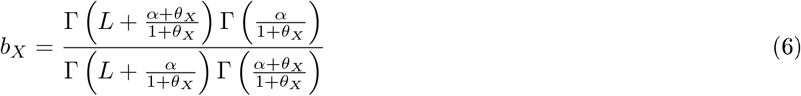

In Appendix B (Supp. Text), we show that, in the limit of very tight linkage between barrier loci (for which we expect the deleterious block to act as a single locus with net effect *S*_*X*_ = *s*_*X*_*L*), our results for barrier strength are consistent with those of Muirhead and Presgraves (2016), who consider a neutral marker linked to a *single* incompatible locus.

### Model extensions

An important feature of the above equations is that any kind of sex bias can be encapsulated by the composite parameters 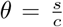 and 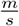, at least for weak migration, selection and recombination. Thus equations (3) and (4) describe X-linked introgression in various scenarios.

#### Dosage compensation and sex-biased forces

Dosage compensation is likely to influence fitness effects of incompatibilities in complex ways that are poorly understood. For simplicity, we focus on two different fitness models, which represent two extreme possibilities. We first analyze a “basic model” with co-dominant alleles in which heterozygous selective effects in females are assumed to be half the corresponding homozygous effects and equal to the hemizygous selective effects in males (i.e., *s*_*M*_ = *s*_*F*_ = *s*_*hom,F*_/2). This parameter setting for our “basic model” is justified to the extent that we first want to focus solely on the effects of lower effective recombination on the sex chromosomes. Moreover, it gives the most conservative estimate of how much barrier strength is increased on the sex chromosome relative to autosomes, because stronger deleterious effects in males (relative to females) further enhance the strength of coupling (*θ*) on the X relative to autosomes (if *s*_*F*_ and *c* are comparable for the two chromosome types).

We then consider a more realistic parameter setting, where we explicitly account for the fact that a single deleterious allele is (typically) more detrimental in the heterogametic sex than a single deleterious allele in the homogametic sex. This setting follows from a model with dosage compensation, where the fitness of a male carrying a hemizygous XY deleterious block is the same as that of a female carrying a homozygous XX block (i.e., *s*_*M*_ = *s*_*hom,F*_ > *s*_*F*_ for sex-linked alleles). Under this assumption, the selection coefficient in males, *s*_*M*_, is larger in magnitude than in the “basic model”, due to the larger amounts of deleterious gene products in the heterogametic sex. This model can describe any mechanism of dosage compensation which leads to similar total levels of activity of the sex chromosome in both sexes. It is thus reasonable for groups in which dosage compensation evolved by up-regulation of the X (or Z) in the homogametic sex. But it can also be applied to groups in which the X (or Z) is initially over-expressed in both sexes, then secondarily down-regulated in the homogametic sex. Note that the assumptions about the influence of dosage compensation on selective effects can be easily relaxed; our mathematical results are very general and can be directly applied to any model of dosage compensation (which would result in different *s*_*F*_/*s*_*hom,F*_ and *s*_*F*_/*s*_*M*_ ratios).

We then model sex-specific selection on deleterious alleles by assuming stronger selection coefficients on XY males (i.e., *s*_*M*_ > *s*_*F*_) or on ZZ males (i.e., *s*_*F*_ > *s*_*M*_ in our XY notation). We study sex-biased migration by setting *m*_*M*_ ≠ *m*_*F*_, and achiasmy by setting *c*_*M*_ = 0 on autosomes in males (or *c*_*F*_ = 0 in a ZW system), such that there is no recombination on any chromosome in the heterogametic sex.

#### Beyond the weak selection approximation

The analytical predictions presented so far assume that selection against hybrids is weak, i.e., *S* = *sL* ≪ 1, (for a given *θ*, *m/S*, and *L*). Under weak selection, the frequency of hemizygous males (carrying a particular X-linked deleterious block) is found to be half the corresponding frequency of heterozygous females, i.e., *P*_*M*_*(y)* ~ *P*_*F*_*(y)*/2. This forms the basis of a relatively simple description of X-linked introgression in terms of sex-averaged X-linked parameters (equations (4) and (6)).

Under stronger selection, homozygous female immigrants may have strongly reduced fitness as compared to hemizygous males: this reduces the relative contribution of females to the next generation, increasing the ratio *P*_*M*_*(y)*/*P*_*F*_*(y)* above 1/2. Moreover, this effect is sensitive to the dominance coefficient of deleterious alleles *(h)*, which determines the relative fitness of homozygous and heterozygous females. However, even under these scenarios (i.e. strong selection and dominance/recessivity), male and female genotypic frequencies satisfy a set of linear equations, as long as the frequency of deleterious alleles in the recipient species is sufficiently low that second order terms in *P(y)* can be neglected. In Appendices A and B (Supp. Text), we present these more general predictions for the equilibrium frequencies *P*_*F*_*(y)* and *P*_*M*_*(y)* as well as for the barrier strengths, for both autosomal and X-linked introgression. These are more accurate when selection against hybrids is strong (see Results), but they are considerably more involved than the weak-selection expressions in eqs. (4) and (6), and cannot be expressed in terms of sex-averaged effective parameters (i.e. *s*_*X*_, *m*_*X*_, *c*_*X*_).

### Individual-based simulations

The analytical expressions for the deleterious allele frequency (eq. (4) which is valid for *sL*, *cL* and *m* ≪ 1; eq. (16) in Appendix A which is also valid for strong selection) and for the barrier strength (eq. (6); eq. (30) in Appendix B) assume that: (i) drift is negligible (*Ns* ≫ 1); (ii) multiple crossovers are rare (*cL* ≪ 1); and (iii) introgressed alleles are rare enough that most individuals carry at most one introgressed block. The last assumption is the most critical, as its validity depends on the value of *θ*. Therefore, we compare our analytical results to forward-in-time individual-based simulations (see Supp. Text for more details) with *Ns* ≫ 1 and *cL* ≪ 1, but in different coupling regimes, for which assumption (iii) may not hold.

## RESULTS

Extending previous results (Barton 1983; Barton and Bengtsson 1986) to the case of sex chromosomes, we have derived analytical expressions for the equilibrium frequency of multilocus sex-linked incompatibilities, and their strength as barriers against gene exchange between two hybridizing species (eqs. (4) and (6), see also Appendices A and B). We now focus on the biological consequences of there being different sex-averaged effective parameters (Table 2 and SI). We summarize the influence of (i) the genetic architecture of the barrier between species, (ii) sex-biased forces and dosage compensation, and (iii) other model extensions (dominance and epistasis, see Supp. Text). We restrict attention to the effect of each factor separately, although combined effects are discussed for the interpretation of empirical data (see Discussion). To be consistent with the mathematical simplifications in our model (see Model and Methods), we choose *sL* to be in the range 0.01 to 0.1 *cL* to be in the range 0.025 to 0.25 and migration to be weak relative to selection coefficients (*m* = 0.001). In simulations, we take population sizes to be sufficiently large (*N* = 10^5^) to conform with the deterministic regime.

### Genetic architecture of the barrier between species

#### Effects of coupling between barrier loci, and of the migration-to-selection ratio

Validating Barton (1983)’s autosomal results, we find that the equilibrium frequency of deleterious alleles decreases as coupling between selected loci becomes stronger, i.e., as *θ* increases (Figure 1A). When coupling between barrier loci is strong (*θ*_*A*_ = 2, red; Figure 1A), the deleterious block acts as the unit of selection, so that the deleterious allele frequency in the recipient species 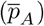 is lower than under intermediate coupling (*θ*_*A*_ = 1, black; Figure 1A). When coupling is weak (*θ*_*A*_ = 0.2, blue; Figure 1A), recombination breaks the genomic block down to smaller sub-blocks of weaker selective disadvantage, resulting in a much higher equilibrium frequency of the introgressing alleles. Note that when *θ* < 1 and *L* is large, the analytical predictions break down: 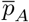 obtained by simulations (open symbols) is much higher than the predicted values (lines and filled symbols). Coupling has a similar effect on gene flow at a neutral marker, i.e. effective neutral migration decreases as *θ* increases (and the barrier strength correspondingly increases; Figure 2), which corroborates Barton and Bengtsson (1986)’s autosomal results. The equilibrium frequency of deleterious alleles in the recipient species shows a strong dependence on 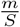 (the ratio of the migration rate to the selective effect *S* = *sL* of the full block) and is in fact equal to 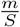 when selected loci are completely linked. Even with incomplete linkage (i.e., strong to intermediate coupling), equilibrium allele frequency is nearly proportional to 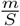: for example, the autosomal equilibrium frequency for low 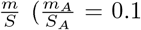, blue; Figure 1B) is ~ 10 times higher relative to that in the high 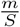 regime (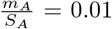, red; Figure IB), nearly independently of the number of barrier loci.

**Figure 1.**
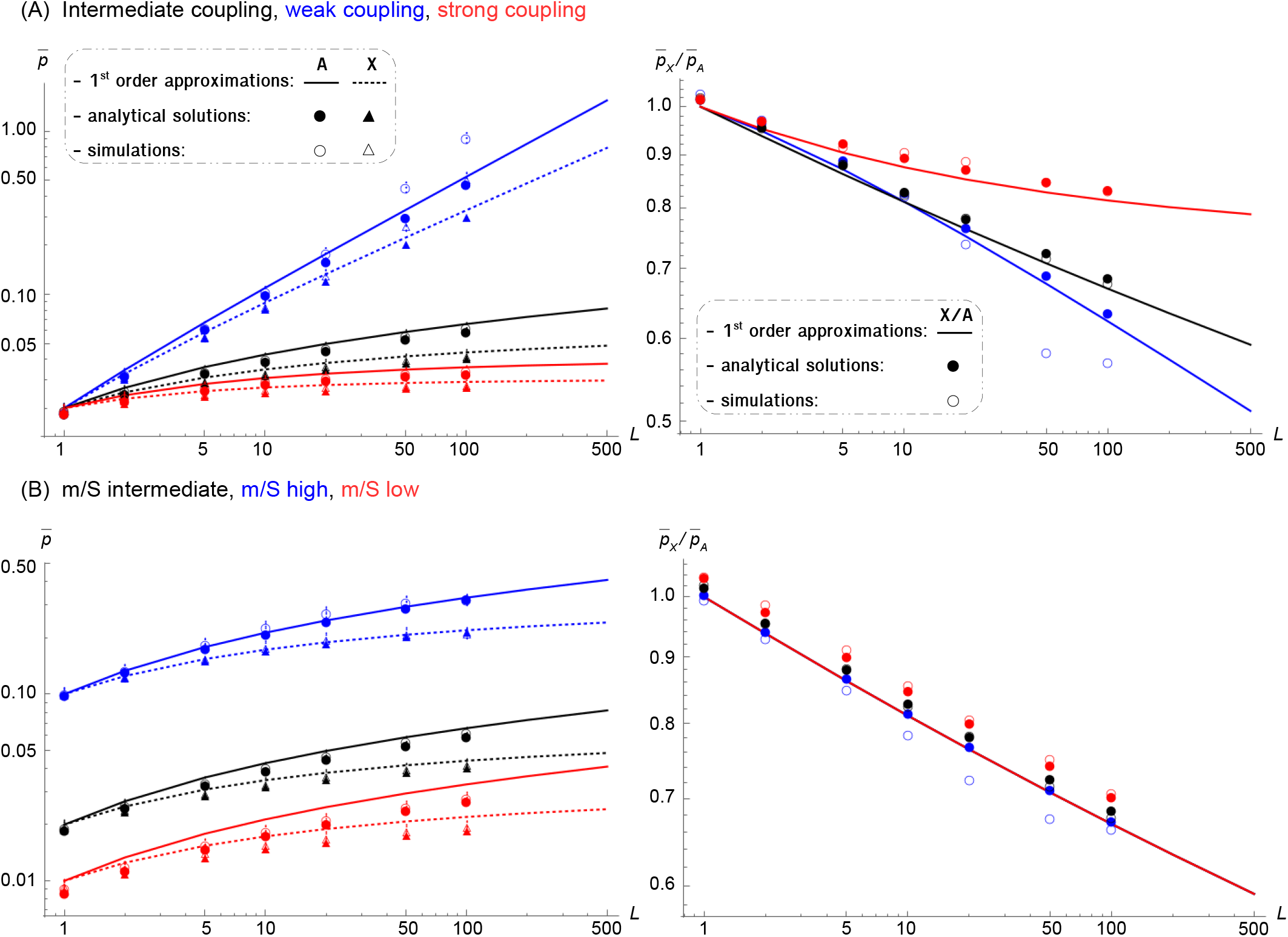
**Effect of *θ* and 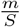 on average equilibrium frequencies** Average frequency of the deleterious alleles in the recipient species (Autosomal: 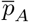; X-linked: 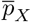; their ratio: 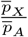) plotted against the number of selected loci on the genomic block, L. The effect of two composite parameters governing introgression is shown: (A) 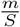 determines the expected allele frequency when selected loci are perfectly linked; (B) 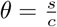 determines the extent of coupling between loci. Lines show the analytical expression in eqs. (4) and (12) (which are given in terms of composite parameters and are valid for *sL cL* and *m* ≪ 1); filled symbols show analytical expression in eqs. (11) and (16) (which are also valid for strong selection); empty symbols show results of individual-based simulations iterated for 10,000 generations and averaged over 100 replicates (error bars indicate the standard error of the mean). Colors stand for values of the composite parameters, *θ* and 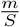. Parameter values for intermediate *θ* and 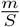 are: *h* = 0.5, *m* = 0.001 *cL* = 0.05, *sL* = 0.05, *m*_*X*_/*S*_*X*_ = *m*_*A*_/*S*_*A*_ = 0.02, *θ*_*X*_ = 1.5, *θ*_*A*_ = 1 and *N* = 10^5^ (simulations). Parameters for weak coupling: *cL* = 0.25, *θ*_*X*_ = 0.3, *θ*_*A*_ = 0.2; strong coupling: *cL* = 0.025, *θ*_*X*_ = 3, *θ*_*A*_ = 2; m/S high: *cL* = *sL* = 0.01, *m*_*X*_/*S*_*X*_ = *m*_*A*_/*S*_*A*_ = 0.1; m/S low: *cL* = *sL* = 0.1 *m*_*X*_/*S*_*X*_ = *m*_*A*_/*S*_*A*_ = 0.01. Note that the total selective disadvantage of the block (*S* = *sL*) and its total map length (*C* = *cL*) are kept constant as we vary the number of loci (L; x axis).

**Figure 2.**
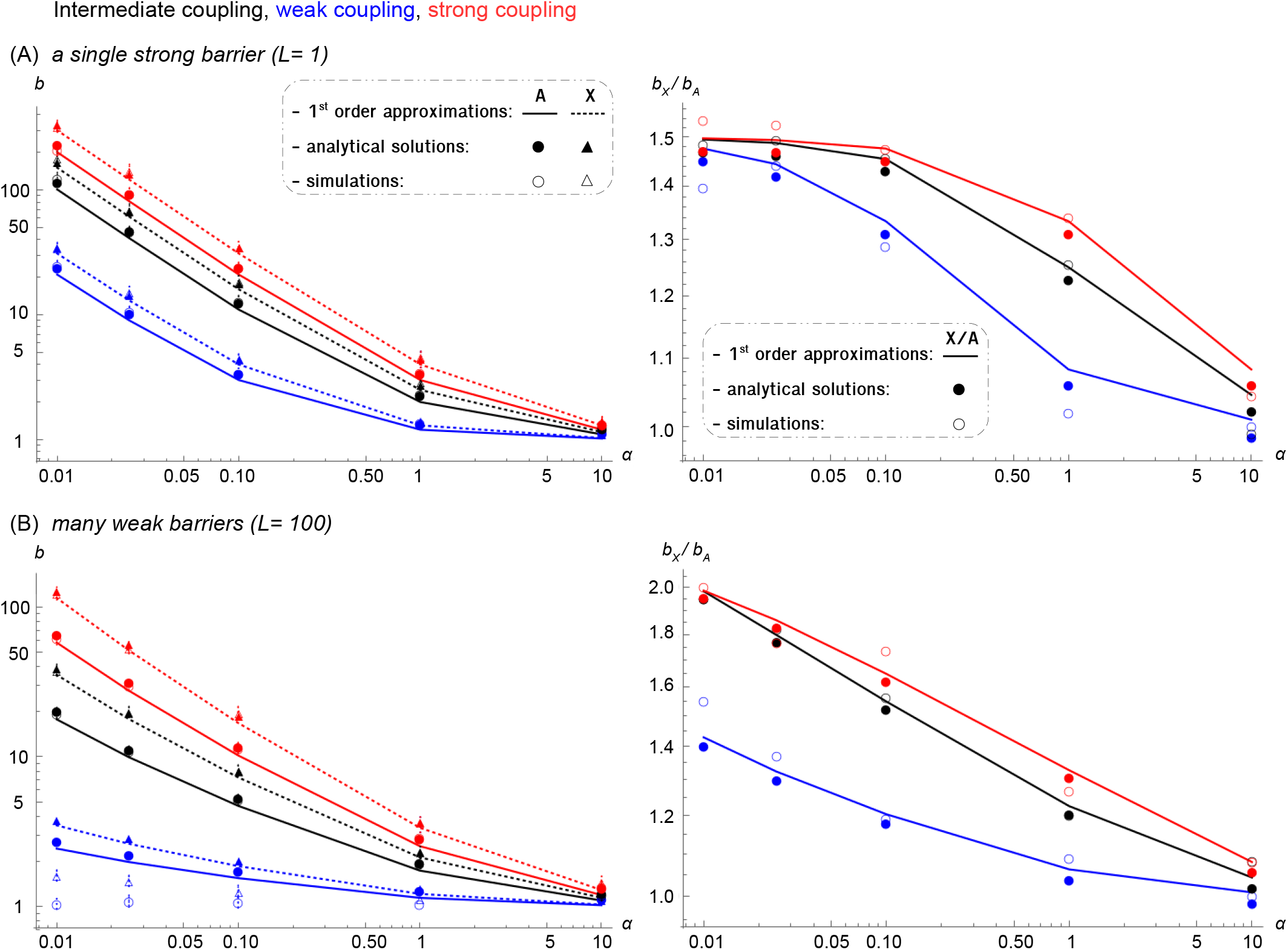
Effect of *θ* on barrier strength at the neutral marker. Barrier strength at the neutral marker (Autosomal: *b*_*A*_; X-linked: *b*_*X*_; their ratio: 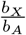) plotted against its proximity to the nearest selected locus, *α*. In (A) the neutral marker is linked to a single strongly selected locus (*L* = 1); while in (B) it is linked to one hundred weakly selected loci (*L* = 100). Lines show the analytical expression in eqs. (5) and (6) (which are given in terms of composite parameters and are valid for *sL, cL* and *m* ≪ 1); filled symbols show analytical expression in eqs. (23) and (30) (which are also valid for strong selection); empty symbols show results of individual-based simulations iterated for 10,000 generations after equilibrating at the selected loci; they are averaged over 100 replicates (error bars indicate the confidence interval of the estimate). Colors stand for values of the composite parameter 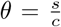 at the selected loci. The recombination rate between the neutral marker and the nearest selected locus is *αLc*, therefore lower *α* values means closer proximity. Other details match Figure 1.

We show that these patterns also hold for the sex chromosomes, but with a consistently lower equilibrium frequency and higher barrier strength (Figure 1A and Figure 2). This is because deleterious alleles on the sex chromosome are more strongly coupled to each other than on autosomes (i.e., 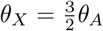; Table SI), while 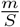 ratios are equal for autosomes and sex chromosomes (i.e., 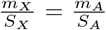; Table SI), assuming identical genetic architectures of hybrid incompatibility for both (i.e. same *s*_*F*_, *s*_*M*_, *c*_*F*_ and *L*). These relationships result in a lower sex-linked equilibrium frequency when the barrier is multilocus (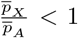 for L > 1; Figure 1A), and a higher barrier strength on the sex chromosome even when there is a single barrier locus (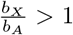 for L ≥ 1; Figure 2), since the neutral marker and the selected locus are then more strongly linked. Note that the difference between X-linked and autosomal introgression probabilities is quite pronounced for low to intermediate values of *θ* (Figure 1A), but becomes weaker for *s*_*F*_, *s*_*M*_ ≫ *c*, i.e., *θ* ≫ 1 (red; Figure S4A), even though X-linked incompatibilities are more strongly coupled than autosomal ones (i.e., 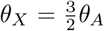) in both regimes. This simply reflects the fact that equilibrium frequencies are nearly independent of the exact value of the coupling strength, when this is large, i.e., when selection acts on the block as a whole.

#### Number of barrier loci on the incompatible block

The dependence of the equilibrium frequency of introgressing alleles on the number of barrier loci, *L*, is qualitatively different in different coupling regimes. In the strong coupling regime (*θ* ≫ 1), the equilibrium frequency of sex-linked and autosomal alleles is largely governed by the net selective disadvantage of the block (*S* = *sL*), and increases only weakly with *L* for fixed *S* (red; Figure 1A and Figure S4A). For smaller *θ*, 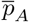 and 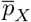 both increase with *L* (black, Figure 1A); correspondingly, the barrier strengths, *b*_*A*_ and *b*_*X*_, decrease (Figure S3). For very weak coupling (*θ* ≪ 1), the equilibrium deleterious allele frequency increases so strongly with *L* that our predictions break down at large *L* (blue; Figure 1A). As before, this is because in the *θ* ≪ 1 regime, any locus is only weakly affected by other selected loci and the effective selection coefficient per allele is close to the raw selective effect *s.* The latter decreases as we consider genetic architectures with larger numbers of more weakly selected loci (from left to right along the x axis in Figure 1).

We also predict that the ratio of autosomal and X-linked deleterious allele frequencies and barrier strengths should increase with the number of barrier loci involved *(L)*, for weak coupling between selected loci. In this regime, the effective strength of selection against individual loci scales as 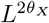 and 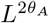 for X-linked and autosomal deleterious alleles respectively; thus, the ratio of equilibrium frequencies scales as 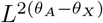 for *θ*_*A*_, *θ*_*X*_ << 1. Since coupling between X-linked alleles is stronger, this means that the ratio of X-linked alleles to autosomal alleles must decrease as the barrier becomes more polygenic. For example, for *θ*_*A*_ = 0.2 and *θ*_*X*_ = 0.3, introgression rates on the sex chromosome are ~ 0.63 times that of the autosomes when the incompatible block is polygenic (*L* = 100, blue; Figure 1A), while, as expected, they are identical for both chromosomes with a single barrier locus (*L* = 1). The same qualitative pattern is observed for the ratio of barrier strengths: 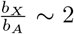 for *L* = 100 while it is ~ 1.5 for *L* = 1, when the neutral marker is close to the block (*α* = 0.01, black; Figure 2). In contrast, under strong coupling (e.g. *θ*_*A*_ = 10 and *θ*_*X*_ = 15), the differences between the two types of chromosomes become negligible (red; Figure S4). Note that for the barrier strength, only the extreme cases of a single strongly deleterious locus (*L* = 1) and many weakly deleterious loci (*L* = 100) are considered hereafter.

#### Distance of the neutral marker from the incompatible block

We have modeled a situation in which a neutral marker lies at one end of an incompatible genomic block (“rod” configuration). The barrier strength always decreases as a function of the distance between the neutral marker and the closest selected locus on both autosomes and the sex chromosomes. This is because the ability of a neutral marker to escape from its incompatible genomic background increases with its map distance from the selected block. For example, under intermediate coupling (black; Figure 2), a multilocus autosomal barrier can reduce the effective migration rate by a factor of ~ 20 for a neutral marker at a relative distance of *α* = 0.01 from the closest barrier locus, while this factor drops to less than ~ 2 at a fraction distance *α* = 1; these numbers are respectively ~ 35 and ~ 2.5 in the sex chromosome case.

This pattern also holds in the more general case of a neutral marker embedded in the selected block (“embedded” configuration, Figure S7). In agreement with previous work on autosomes (Barton and Bengtsson 1986) and sex chromosomes (Muirhead and Presgraves 2016), the barrier at the center is up to one order of magnitude stronger than the barrier at the edge of the block, because two recombination events (instead of one) are required for the neutral marker to escape from the deleterious background. Moreover, as the predictions concerning sex-linked barriers can again be obtained from the corresponding autosomal predictions by a change in sex-averaged effective parameters, our general conclusion that these impede neutral flow more strongly than autosomal barriers still holds in the ‘embedded’ simulations. However, this effect is more extreme than in the “rod” configuration (e.g. 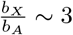 instead of ~ 2 for *L* = 100 and *α* = 0.01; Figure S7).

### Dosage compensation and sex-biased forces

#### Dosage compensation

The basic model considered so far assumes that the hemizygous selective effect of a deleterious allele in a XY male (or a ZW female) is half as strong as the homozygous effect in a XX female (or a ZZ male). However, various molecular mechanisms of dosage compensation have evolved independently in various clades in response to these imbalances (Gu and Walters 2017). These mechanisms may boil down to a fitness model where the effect of a single copy of a sex-linked allele in hemizygous males is similar to that of two copies in homozygous females (*s*_*M*_ = *s*_*hom,F*_ = 2*s* and *s*_*F*_ = *s* for *h* = 0.5). In such a situation, we have *θ*_*X*_ = 2*θ*_*A*_ and 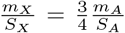 (Table SI), which results in a lower X-to-autosome ratio of introgressed allele frequencies for weak to intermediate coupling between selected loci (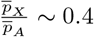 instead of ~ 0.66 in the basic model with *L* = 100, compare red vs. black; Figure 3A). This holds for single and multilocus barriers (i.e., for L ≥ 1), since not only coupling between sex-linked alleles is stronger than in the basic model, but also *m/S* on the sex chromosomes is weaker than on autosomes. Likewise, the influence of dosage compensation on neutral gene flow is to increase the ratio of barrier strengths relative to the basic model (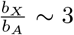 instead of ~ 2 with *L* = 100 and *α* = 0.01, compare red vs. black; Figure 4A).

**Figure 3.**
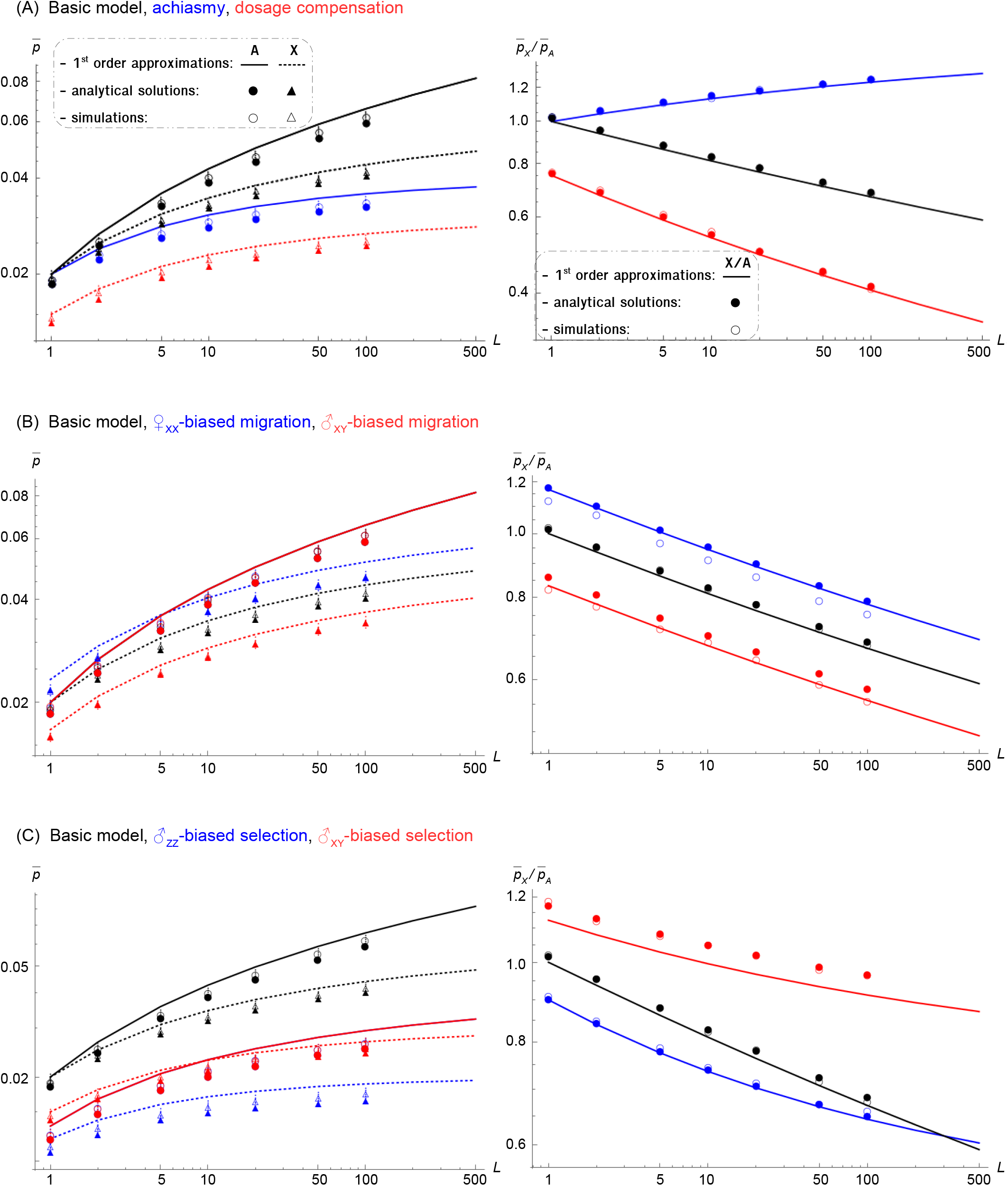
Effect of sex-specificities on average equilibrium frequencies. Average frequency of the deleterious alleles in the recipient species (Autosomal: 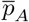; X-linked: 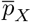; their ratio: 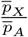) plotted against the number of selected loci on the genomic block, *L*. (A) Dosage compensation (*s*_*M*_*L* = 2*sF*_*L*_ = *s*_*hom,F*_*L* = 0.1 for sex-linked alleles, *m*_*X*_/*S*_*X*_ = 0.015, *θ*_*X*_ = 2, red) and achiasmy in XY males (*c*_*M*_*L* = 0, *θ*_*A*_ = 2, blue). (B) Sex-biased migration: e.g., if males migrate three times more than females (*m*_*M*_ = 3*m*_*F*_ = 0.0015. *m*_*X*_/*S*_*X*_ ~ 0.016, red); or females migrate three times more than males (*m*_*F*_ = 3*m*_*M*_ = 0.0015, *m*_*X*_/*S*_*X*_ ~ 0.023, blue). (C) Sex-biased selection: e.g. if selection coefficient is twice as strong on XY males as on females (*s*_*M*_*L* = *2s*_*F*_*L* = 0.1 *m*_*X*_/*S*_*X*_ ~ 0.015, *m*_*A*_/*S*_*A*_ ~ 0.013, *θ*_*X*_ = 2, *θ*_*A*_ = 1.5, red); or selection coefficient is twice as strong on ZZ males as on females (*s*_*F*_*L* = *2s*_*M*_*L* = 0.1 in our XY notation, *m*_*Z*_/*S*_*Z*_ ~ 0.012, *m*_*A*_/*S*_*A*_ ~ 0.013, *θ*_*Z*_ = 2.5, *θ*_*A*_ = 1.5, blue). The basic model is shown in black in all panels. Parameter values for the basic model are: *h* = 0.5, *m* = 0.001 *cL* = 0.05, *sL* = 0.05, *m*_*X*_/*S*_*X*_ = *m*_*A*_/*S*_*A*_ = 0.02, *θ*_*X*_ = 1.5, *θ*_*A*_ = 1 and *N* = 10^5^ (simulations). Other details match Figure 1.

**Figure 4.**
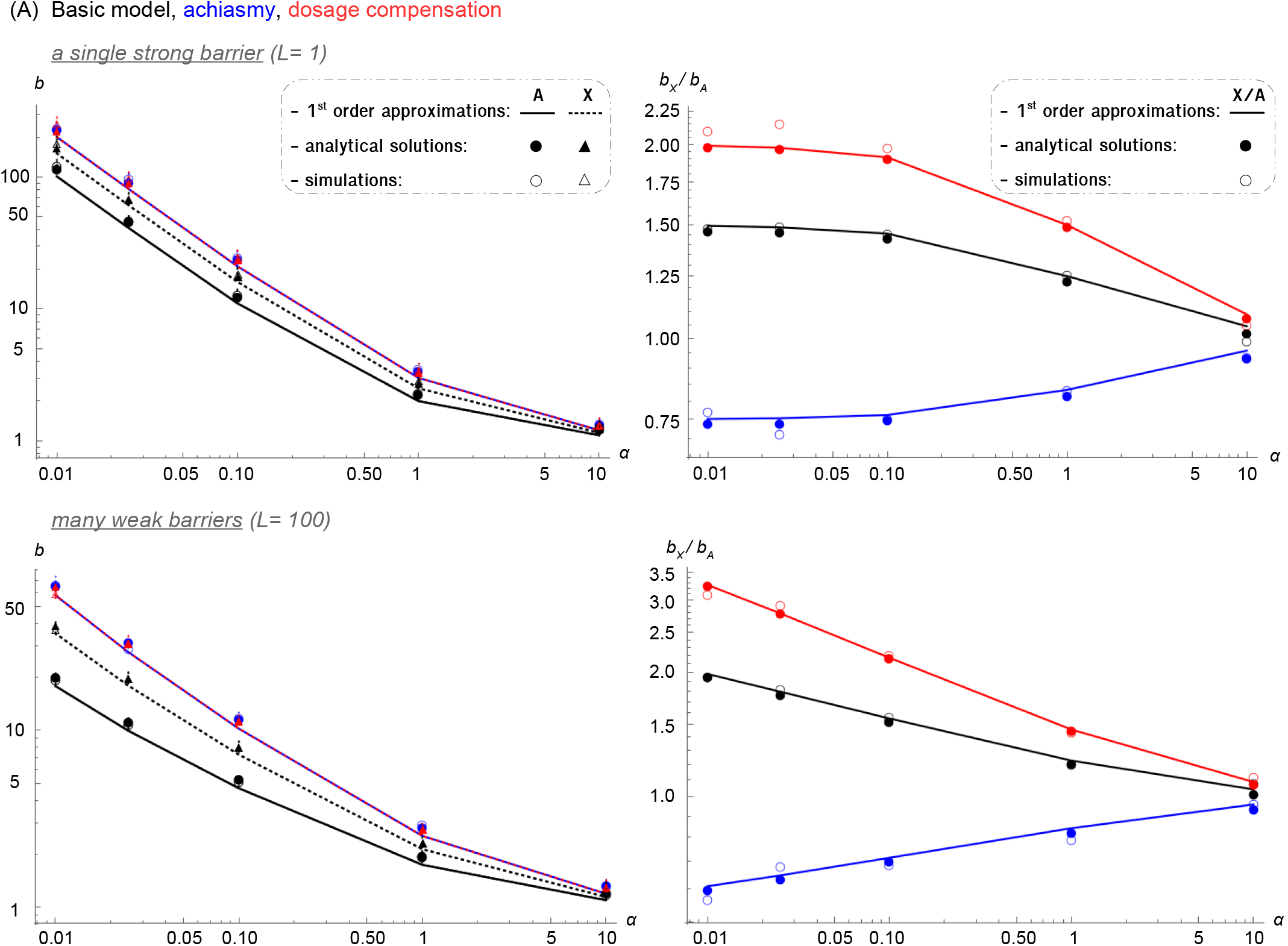
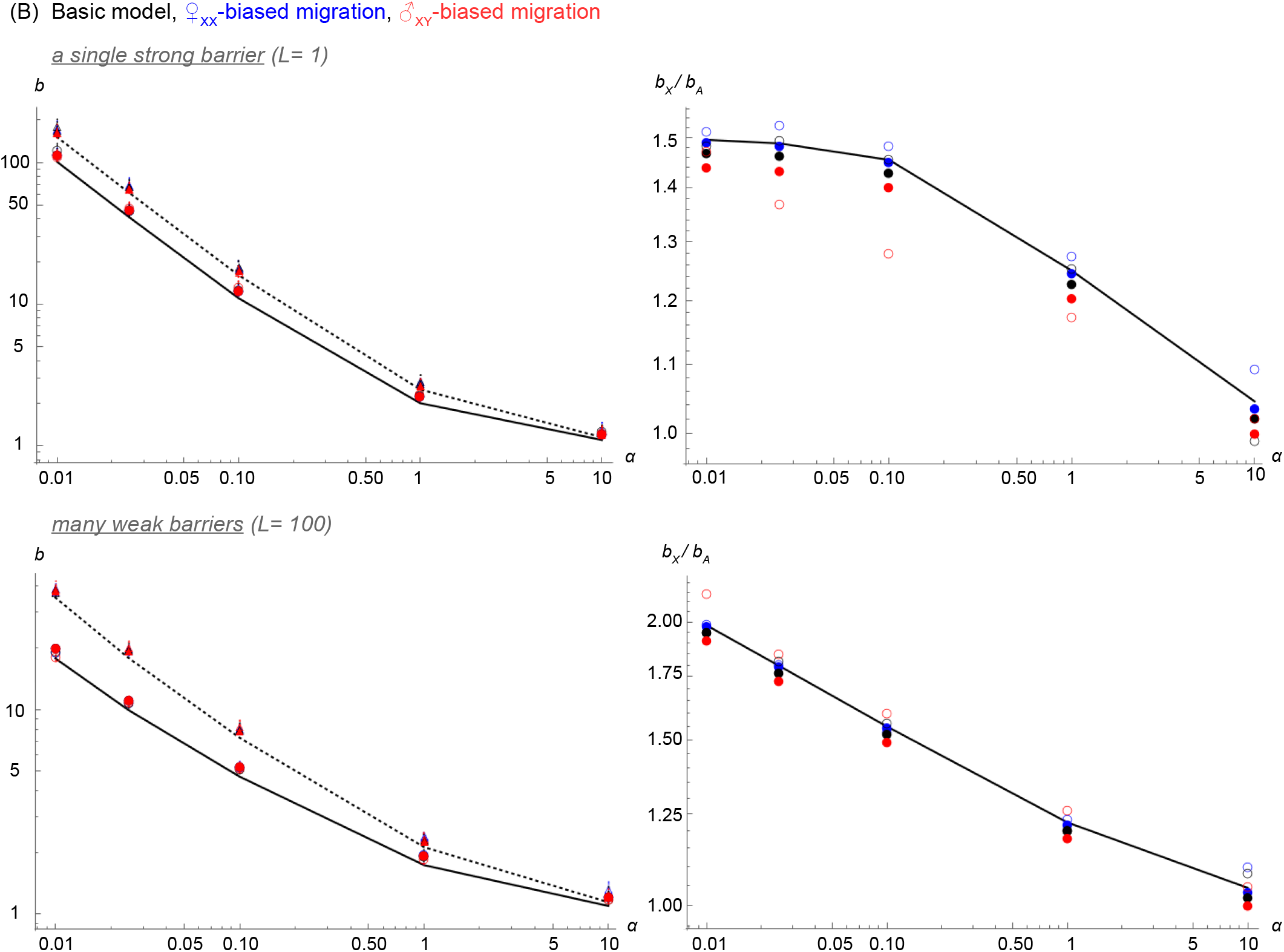
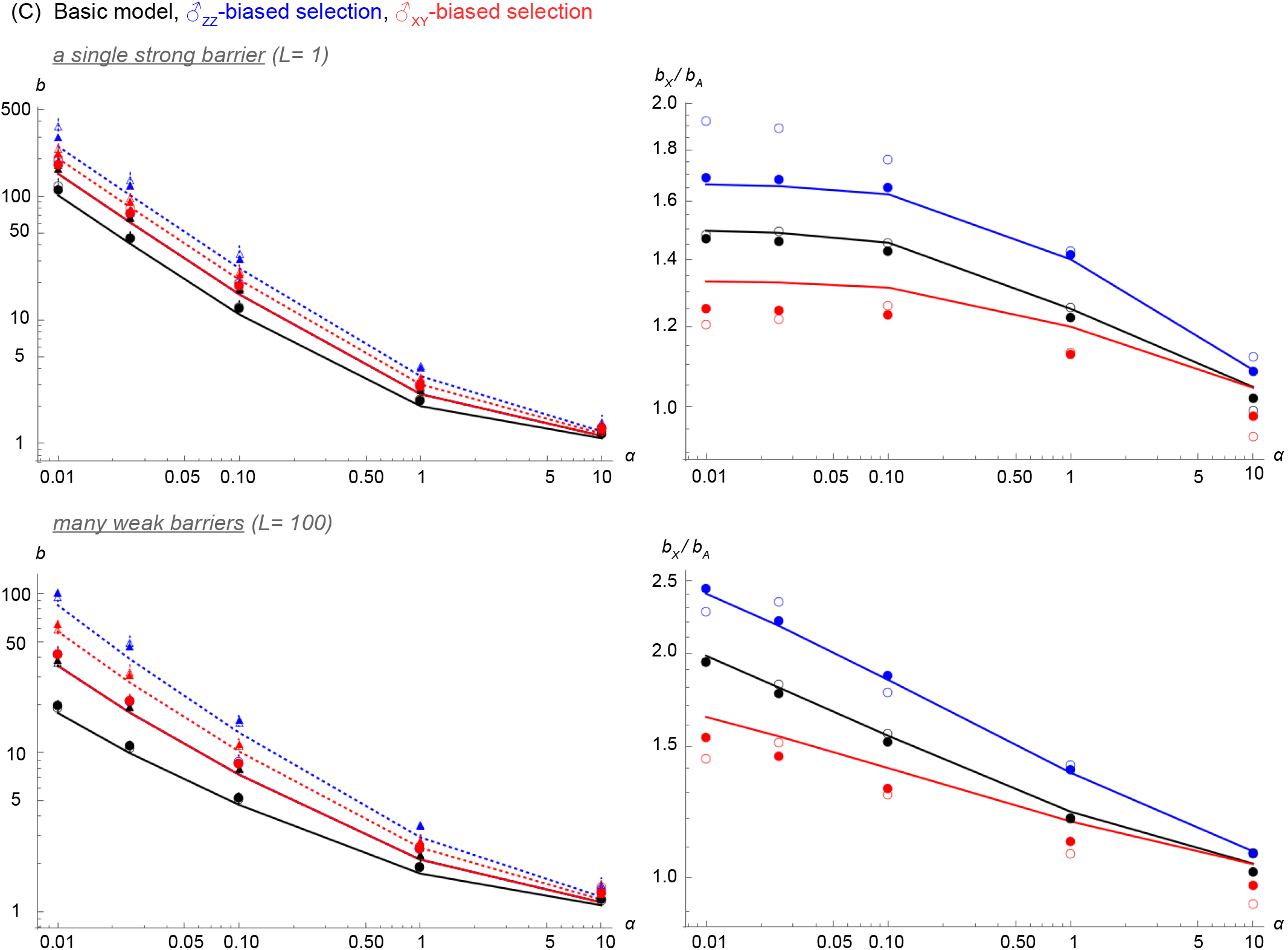
Effect of sex-specificities on barrier strength at the neutral marker. Barrier strength at the neutral marker (Autosomal: *b*_*A*_; X-linked: *b*_*X*_; their ratio: 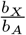) plotted against its proximity to the nearest selected locus, *α*. **(A)**Dosage compensation (red) and achiasmy in XY males (blue). **(B)**Sex-biased migration: e.g., if males migrate three times more than females (red); or females migrate three times more than males (blue). **(C)**Sex-biased selection: e.g. if selection coefficient is twice as strong on XY males as on females (red); or selection coefficient is twice as strong on ZZ males as on females (blue). The basic model is shown in black in all panels. Other details match Figure 2 and 3.

#### Achiasmy

In some species, such as certain dipterans or lepidopterans, recombination is absent throughout the genome in the heterogametic sex (i.e. their meiosis is achiasmate, so *c*_*M*_ = 0 on autosomes and sex chromosomes in an XY sexual system; Satomura et al. 2019). As sex-linked introgressing alleles (whether on the X or Z) spend one third of their time in the heterogametic sex, while autosomal alleles spend an equal amount of time in both sexes (at least under weak selection), achiasmy leads to a lower effective recombination rate of autosomal compared to sex-linked incompatibility loci (all else being equal; Langley et al. 1988, Betancourt et al. 2004). In this case, we have 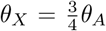 (Table SI), i.e. the coupling between barrier loci is stronger on autosomes than on sex chromosomes. As a consequence, achiasmatic species exhibit higher introgression of sex chromosomes than autosomes when barriers are multilocus (~ 1.2 times higher with *L* = 100, blue; Figure 3A), and their barrier strength is accordingly weaker (~ 0.6 that of autosomes with *L* = 100 and *α* = 0.01, blue; Figure 4A).

#### Sex-biased migration

Differences in migration rates between sexes are commonly observed in nature, with the heterogametic sex prone to higher migration in mammals and birds (Greenwood 1980; Trochet et al. 2016), although there are many exceptions and patterns are less known in other taxa. Sex-biased migration affects only the introgression of the sex-linked alleles (to lowest order in *sL* and *cL*), because immigrants of the heterogametic and homogametic sex introduce different numbers of sex-linked blocks, but the same number of autosomal blocks. Under male-biased migration in a XY system (or female-biased in a ZW system), the migration-to-selection ratio is thus lower for sex-linked alleles than for autosomal alleles (e.g. if *m*_*M*_ = 3*mp*, then 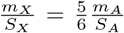 Table SI). Compared to the basic model, this results in a lower X-to-autosome ratio for *L* ≥ 1 and weak to intermediate coupling (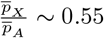 instead of ~ 0.66 with *L* = 100, red; Figure 3B). Note that the barrier strength is predicted to be insensitive to sex-bias in migration to lowest order in *sL* and *cL* (see eqs. (5) and (6)). However, for stronger selection, sex-biased migration can have a very weak effect on barrier strength, which is predicted by eqs. (22) and (30) (Appendix B, and Figure 4B). Opposite patterns are found when migration is female-biased in a XY system (e.g. if *m*_*F*_ = 3*m*_*M*_, then 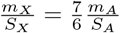; Table SI), or male-biased in a ZW system.

#### Sex-biased selection

In most species, the strength of sexual selection is stronger in males than in females (Singh and Punzalan 2018). Since twice as many females carry a single copy of any X-linked introgressed allele as males (i.e., any X-linked incompatible allele spends twice as much time in females as in males), while the reverse is true for the Z chromosome, male-biased sexual selection will lead to opposite effects in the two sexual systems. For example, if selection is twice as strong on XY males as on females (*s*_*M*_ = 2*s*_*F*_), we have 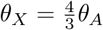 and 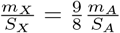 (Table SI), i.e. weaker coupling between barrier loci and a stronger migration-to-selection ratio on the sex chromosome compared to autosomes, than in the basic model. This results in a higher X-to-autosome equilibrium frequency compared to the basic model (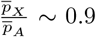 instead of ~ 0.66 with *L* = 100, red; Figure 3C), and a correspondingly lower ratio of barrier strengths (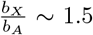 instead of ~ 2 with *L* = 100 and *α* = 0.01, red; Figure 4C). On the contrary, when ZZ males are under stronger selection than females (*s*_*F*_ = 2*s*_*M*_, using our XY notation), we have 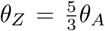 and 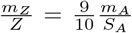 (Table SI), resulting in lower Z-to-autosome equilibrium frequencies compared to the basic model (Figure 3C) and a correspondingly higher barrier strength (Figure 4C).

### Dominance of barrier loci

In Figures 1–4, we have shown results for individual-based simulations in which incompatible mutations have co-dominant effects (*h* = 0.5), and compared these with our analytical predictions, which are actually independent of dominance as long as selection is weak, i.e., *sL* and *cL* ≪ 1 (see Table SI and Appendix A). We now examine how dominance affects introgression patterns by considering either partial recessivity (*h* = 0.1) or full dominance (*h* = 1.0) of the incompatible effects, and by varying their homozygous deleterious effect (*s*_*hom*_) while keeping the heterozygous selective effect *(s)* constant. This allows us to test the influence of *h* = *s/s*_*hom*_ (i.e., the ratio of heterozygous to homozygous effects) for a given heterozygous effect.

Recessivity of deleterious alleles implies that homozygous blocks are far more deleterious than in the co-dominant model (*s*_*hom*_ = 0.5 for *h* = 0.1 vs *s*_*hom*_ = 0.1 for *h* = 0.5), resulting in a strong selective disadvantage for first generation immigrants of the homogametic sex (in case of sex-linked blocks) or of both sexes (in case of the autosomal blocks). This leads to lower levels of introgression for both the sex chromosome and autosomes (blue; Figure S5B). Moreover, total migration is more strongly reduced in autosomes (due to the reduced fitness in both male and female migrants) relative to sex chromosomes (since only female migrants have substantially lower fitness than in the co-dominant model). Thus, the X-to-autosome introgression ratio is higher than that in the co-dominant case (Figure S5B), and the ratio of barrier strengths is correspondingly smaller (Figure S6B). However, it is worth noting that introgression remains low overall in our model, and so homozygous blocks are rare in the population. They are actually neglected in the weak selection approximation (lines) and only enter into the more general analytical predictions (filled symbols) via the fitness of homozygous migrants (of the homogametic sex); however, they are fully considered in the simulations (empty symbols). Therefore, the influence of dominance *(h)* on equilibrium frequencies and barrier strengths is by itself weak: what matters to the first order is only the heterozygous effects *(s)* of the incompatible alleles.

If we also assume dosage compensation, then the migration-to-selection ratio 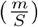 of X-linked recessive alleles is only 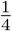 that of the autosomes (compared to 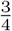 when alleles are co-dominant, see Table SI), and coupling between incompatible alleles (*θ*) is 6 times stronger on the X chromosome relative to autosomes (compared to 2 times when alleles are codominant, see Table SI). Importantly, these two effects cause a much lower X-to-autosome introgression ratio than under co-dominance (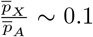 instead of ~ 0.4 with *L* = 100, blue; Figure S5C), and also a much greater X-to-autosome ratio of barrier strengths (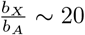 instead of ~ 3.5 with *L* = 100 and *α* = 0.01, blue; Figure S6C). This point in the parameter space is important to consider as it corresponds to very different X-linked effects in hemizygous males and heterozygous females, which is the basis of the dominance theory (Turelli and Orr 1995, 2000).

Under complete dominance of incompatible alleles, homozygous blocks are just as deleterious as heterozygous blocks (*s*_*hom*_ = 0.05 for *h* = 1.0). This leads to slightly higher levels of introgression for both chromosome types in the simulations (red; Figure S5B), compared to the co-dominant model. With dosage compensation, the relationship between the composite parameters is 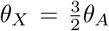 and 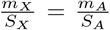 (Table SI); therefore dominant X-linked alleles introgress more than co-dominant alleles (red; Figure S5C), and act as weaker barrier to gene flow (red; Figure S6C).

## DISCUSSION

Previous work has considered the effect of divergence at large numbers of autosomal foci on genetic exchange between hybridizing species (Barton 1983; Barton and Bengtsson 1986; Baird 1995). Here, we have extended the analysis to sex chromosomes (X or Z). Under weak selection, predictions for the introgression of multilocus deleterious blocks (eq. 4), and their strength as barriers to neutral gene flow (eq. 6), are governed by the same equations as for autosomes, (eqs. 12 and 5), but with different sex-averaged effective parameters *m*_*X*_, *s*_*X*_ and *c*_*X*_. We show that all sex-linked effective parameters have the form 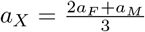, i.e., they are weighted sums of female and male components, *α*_*F*_ and *α*_*M*_. This implies that evolutionary forces that affect males and females differently can be readily encapsulated by these effective parameters, and their biological consequences predicted. An extreme example is that of the X-linked recombination rate, *c*_*X*_, which is 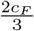, since males do not contribute at all to recombination on the X.

These expressions for sex-linked effective parameters are identical to those proposed by Hedrick (2007) (see also Haldane 1926; Avery 1984), based on the argument that females carry two copies of the X chromosome (versus a single copy in males) and thus are associated with a weight that is twice that of males. However, the reason underlying the 2:1 female and male contributions to *s*_*X*_ and *c*_*X*_ in our model is somewhat different, since both males and females, with the exception of first-generation female immigrants, typically carry a single copy of any X-linked introgressed allele (assuming low-introgression). As we demonstrate above, X-linked alleles nevertheless spend twice as much time in females as in males because twice as many females are heterozygous for X-linked deleterious alleles as males (are hemizygous). However, very strong selection against hybrids elevates the ratio *P*_*M*_*(y)*/*P*_*F*_*(y)* above 1/2 (see equation 15b, Appendix A), causing this simple description in terms of sex-averaged X-linked parameters to break down: analytical predictions can still be derived by separately considering genotypic frequencies in males and females.

### Conditions for an oversized role of sex chromosomes in speciation

We find that the stronger coupling of incompatible alleles on the sex chromosome reduces the escape probability of neutral markers, and thus enhances their barrier strength by a factor of ~ 1.5 to ~ 3 relative to autosomes under optimal conditions (i.e. when the neutral marker is close to a multilocus barrier). This result mirrors earlier predictions for single and two-locus barriers under a regime where introgressed alleles are rare (Muirhead and Presgraves 2016), and for the maintenance of DMIs in parapatry (Hoellinger and Hermisson 2017). Although it is a moderate effect, this process is very general as it applies to species diverging in parapatry under various scenarios (including dosage compensation, sex-biased migration and sex-biased selection; see below), and regardless of the position of the neutral marker along the chromosome. However it may not be sufficient to explain the major role of sex chromosomes in the speciation of achiasmatic species such as *Drosophila*, where an opposite effect is expected (i.e. X-linked barriers are weaker than autosomal ones) if achiasmy is the sole sex-biased process acting.

The greatest sex chromosome-to-autosome ratio for the barrier strengths 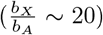 occurs when incompatible alleles are recessive and dosage compensated (Figure S6C), in agreement with the model of Muirhead and Presgraves (2016). In contrast, Hoellinger and Hermisson (2017) did not find a large difference in the outcomes between co-dominant and recessive DMIs. We note that in their model the level of dominance is not associated with the single-locus effects but with the two-locus DMI itself, and so the fitness setting is unlike ours. Importantly, under our model assumptions, dominance (i.e. the ratio of heterozygous to homozygous fitness) itself has little effect as it affects only the selective disadvantage of migrants in the first generation (and of rare homozygous individuals). In essence, recessivity and dosage compensation cause hemizygous deleterious effects of X-linked alleles in males to be much stronger than the corresponding heterozygous effects in females. This is in line with the dominance theory (Turelli and Orr 1995, 2000), which proposes that if most incompatible alleles act recessively in hybrids, then their deleterious effects would be more exposed on the sex chromosomes (as they are hemizygous in males). Therefore, selection more efficiently removes recessive incompatible alleles from the sex chromosome, and eventually imposes a stronger barrier to neutral flow.

A new prediction that emerges from our multilocus model is that polygenic barriers comprised of many weakly coupled alleles result in a greater sex chromosome-to-autosome difference than strong single-locus barriers (although the absolute strength of the barrier is stronger on the latter case). Therefore, the genetic architecture of reproductive isolation affects the extent of introgression on sex chromosomes relative to autosomes, which may have important implications. QTL mapping studies for sterility and/or inviability of hybrids typically detect a few large-effect loci (Maheshwari and Barbash 2011), but numerous smaller-effect incompatibilities may be commonly underestimated in these studies. Detailed backcrossing experiments have actually shown that each introgressed block might contain multiple linked barrier loci each required for reproductive isolation to appear (for a review see Fraïsse et al. 2014). Such polygenic barriers between species, if common, may contribute to the oversized role of sex chromosomes in speciation.

### The influence of dosage compensation

Dosage compensation mechanisms are diverse among taxa (Gu and Walters 2017). In dipterans and hemipterans, imbalance of the sex-linked gene dose is solved by an up-regulation of the sex chromosome in the heterogametic sex, while in nematodes and mammals, there is a down-regulation of the sex chromosome in the homogametic sex in response to an early over-expression in both sexes (Charlesworth 1978). These evolutionary mechanisms are equivalent in terms of fitness effects as long as they lead to similar total levels of activity of the X (or Z) in males and females. We have shown that dosage compensation (e.g., the X expression is doubled in XY males) reinforces the barrier strength ratio of sex chromosomes to autosomes compared to the basic setting (Figure 4A), consistent with findings of Hoellinger and Hermisson (2017). This is because a single X-linked incompatible allele in males is selected against twice as strongly as a single X-linked allele in (heterozygous) females, while selection against hemizygous males and heterozygous females is equally strong in the basic model.

However, various complications exist. First, there is a controversy over whether there has indeed first been upregulation of the X (or Z) in both sexes in response to degeneration of the Y (or W), followed by down-regulation in the homogametic sex (e.g. see the work by Mahajan and Bachtrog 2015 in *Tribolium).* Second, there are indications that dosage compensation is partial in some clades like in birds (Graves 2014). This means that the average expression of Z-linked dosage-sensitive genes in females is typically below that of the two copies in males. Third, the random inactivation of one of the two X chromosomes in mammalian females makes the situation even more complicated, as females are functionally hemizygous, but at the cell level (except in marsupials where paternal X inactivation makes females phenotypically hemizygous at the individual level, like males; Watson and Demuth 2012). Therefore, the effect of a heterozygous mutation on fitness depends on whether gene expression is cell autonomous (likely causing semi-dominance for fitness, Mank et al. 2010) or not. Finally, although our dosage compensation model assumptions are in line with precedents in the theoretical literature (e.g. Avery 1984), to the best of our knowledge, there is no direct proof that supports these.

### The influence of sex-biased forces

Evolutionary parameters governing introgression and barrier strength (i.e., *m*, *s* and *c*) may differ significantly between the sexes (Hedrick 2007). One general trend is that migration is biased towards the heterogametic sex (males in mammals and females in birds; Greenwood 1980; Trochet et al. 2016). As the heterogametic sex carries only one copy of the X (or Z) vs. two for each autosome, this type of sex-bias leads to a deficit of the number of migrating X (or Z) relative to autosomes. Accordingly, we observed that both male-biased migration (in XY systems) and female-biased migration (in ZW systems) reduce the deleterious equilibrium frequency in the recipient species (Figure 3B, and see Hoellinger and Hermisson 2017 for a similar effect on stability conditions of DMIs in parapatry); thus we do not expect systematic differences between the two sexual systems. Moreover, in the weak selection limit (*sL* ≪ 1), the barrier strength is predicted to be independent of sex-biased migration (see eqs. (5) and (6)); even under stronger selection, the ratio *m*_*F*_/*m*_*M*_ has only an extremely weak effect on *b* (see Figure 4B; eqs. (22) and (30)).

Second, quantitative differences in recombination rate between sexes are common (heterochiasmy; Lenormand and Dutheil 2005). In some cases, recombination is totally absent in the heterogametic sex (Satomura et al. 2019), causing the effects of linkage to be stronger on autosomes relative to the sex chromosome. As sex-linked alleles spend less time in the heterogametic sex (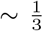 of their time in XY males or in ZW females) than autosomal alleles (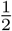 of their time), achiasmy enhances the opportunity for neutral alleles to escape from the sex-linked incompatible loci. As a consequence, incompatibilities on the sex chromosomes become weaker interspecies barriers than autosomes (Figure 4A, qualitatively consistent with the observations of Muirhead and Presgraves 2016). Again, this effect is small and should not produce systematic differences between the X and the Z chromosomes, but is still important to consider in speciation genetics as many detailed studies have been done in *Drosophila*, where achiasmy is widespread.

Third, sexual selection tends to be stronger in males relative to females (Mallet et al. 2011; Sharp and Agrawal 2013; Singh and Punzalan 2018). If sexual selection contributes to reproductive isolation between incipient species, then the impact of male-biased sexual selection should vary between XY systems (where males are heterogametic and an X-linked allele spends 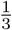 of its time in males) and ZW systems (where males are homogametic and a Z-linked allele spends 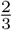 of its time in males). In agreement with this prediction, and assuming a two-fold selection bias toward males, we show that the barrier strength of the Z relative to autosomes is a little increased compared to the basic model, while for the X it is correspondingly decreased (Figure 4C).

### Insights from empirical data

Next-generation data from a diversity of organisms has provided methods to scan genomes for interspecies reproductive barriers, and contrast autosomes with sex chromosomes. Provided that speciation takes place in the presence of gene flow, we expect incompatibilities and their surrounding loci to exhibit a lower rate of interspecific introgression than the rest of the genome (Roux et al. 2013; Sethuraman et al. 2019). In contrast, if speciation occurs in allopatry, it becomes impossible to locate barrier loci, because interspecies divergence is then a simple function of the time elapsed since isolation, which is shared by the whole genome (Wilkinson-Herbots 2008). By putting together population genomic studies of speciation, Presgraves (2018)’s meta-analysis provides useful insights into the role of sex chromosomes (X or Z) in speciation. He shows that sex chromosomes have a systematically higher differentiation level relative to autosomes in 101 species pairs for which *F*_*ST*_ was reported. However, as acknowledged by the author, the impacts on genomic divergence of confounding factors occurring within species (like demographic events or linked selection) are hard to tell apart from the effect of selection against migrants between species, especially when summary statistics are applied.

To address this issue, we report in Table 3 empirical studies that statistically evaluate speciation models from genomic data (at the two extremes: a model of allopatric speciation vs a model where speciation is opposed by continuous gene flow; Sousa and Hey 2013), and then test whether sex chromosomes are more resistant to interspecies introgression than autosomes by estimating the introgression rate for each type of chromosome. Since cline analyses also provide unambiguous estimates of barrier strength (Barton and Hewitt 1985), and simulation studies have shown that incompatible sex-linked alleles flow more slowly than autosomal alleles across a cline (Wang 2013; Hvala et al. 2018; Sciuchetti et al. 2018), we also include them in Table 3, together with studies inferring introgressed tracts from phased genomes (Lawson et al. 2012). We investigate a wide range of species (34 species pairs, including 16 XY systems and 18 ZW systems) characterized by diverse mechanisms of dosage compensation and sex-biased forces.

**Table 3.**
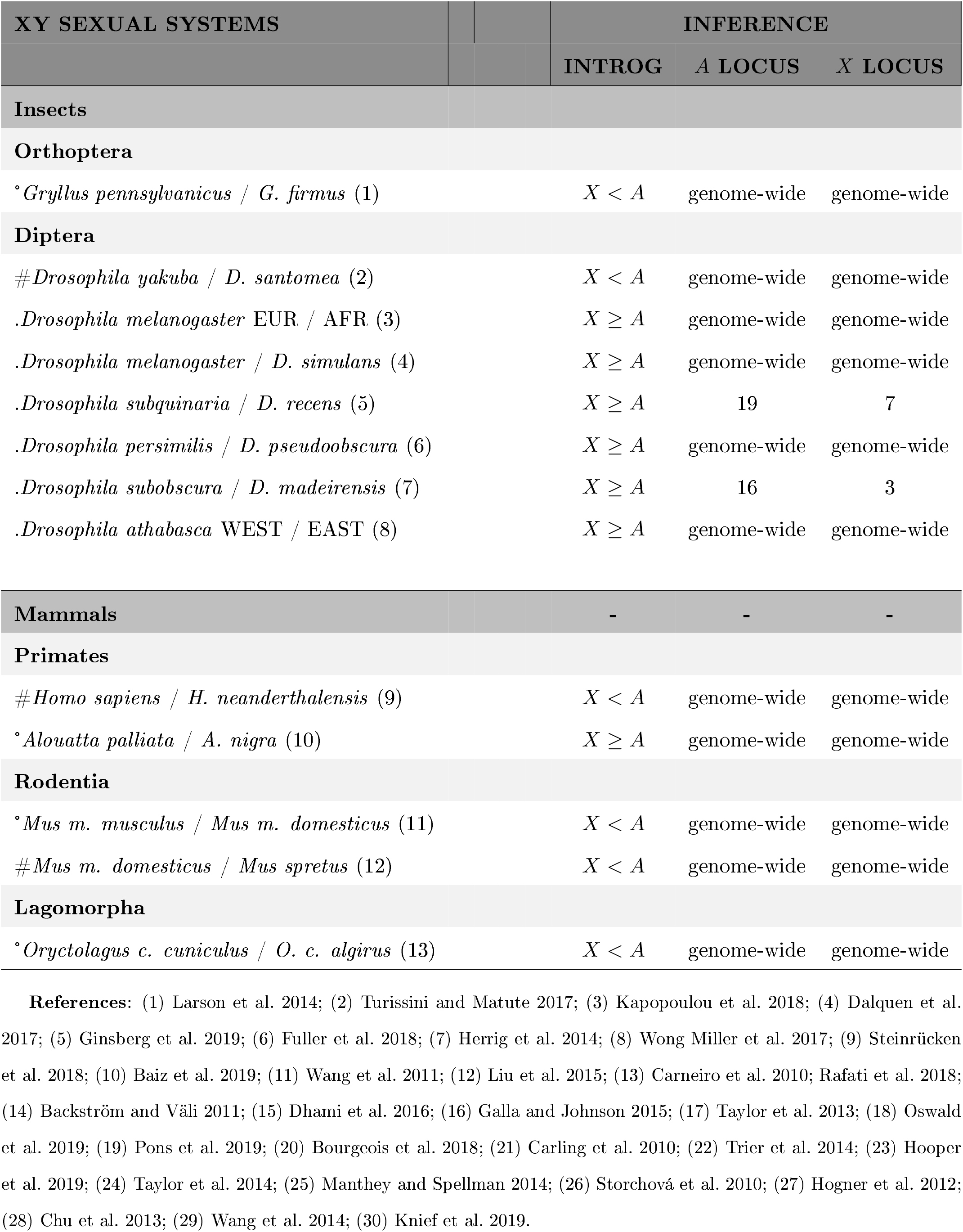
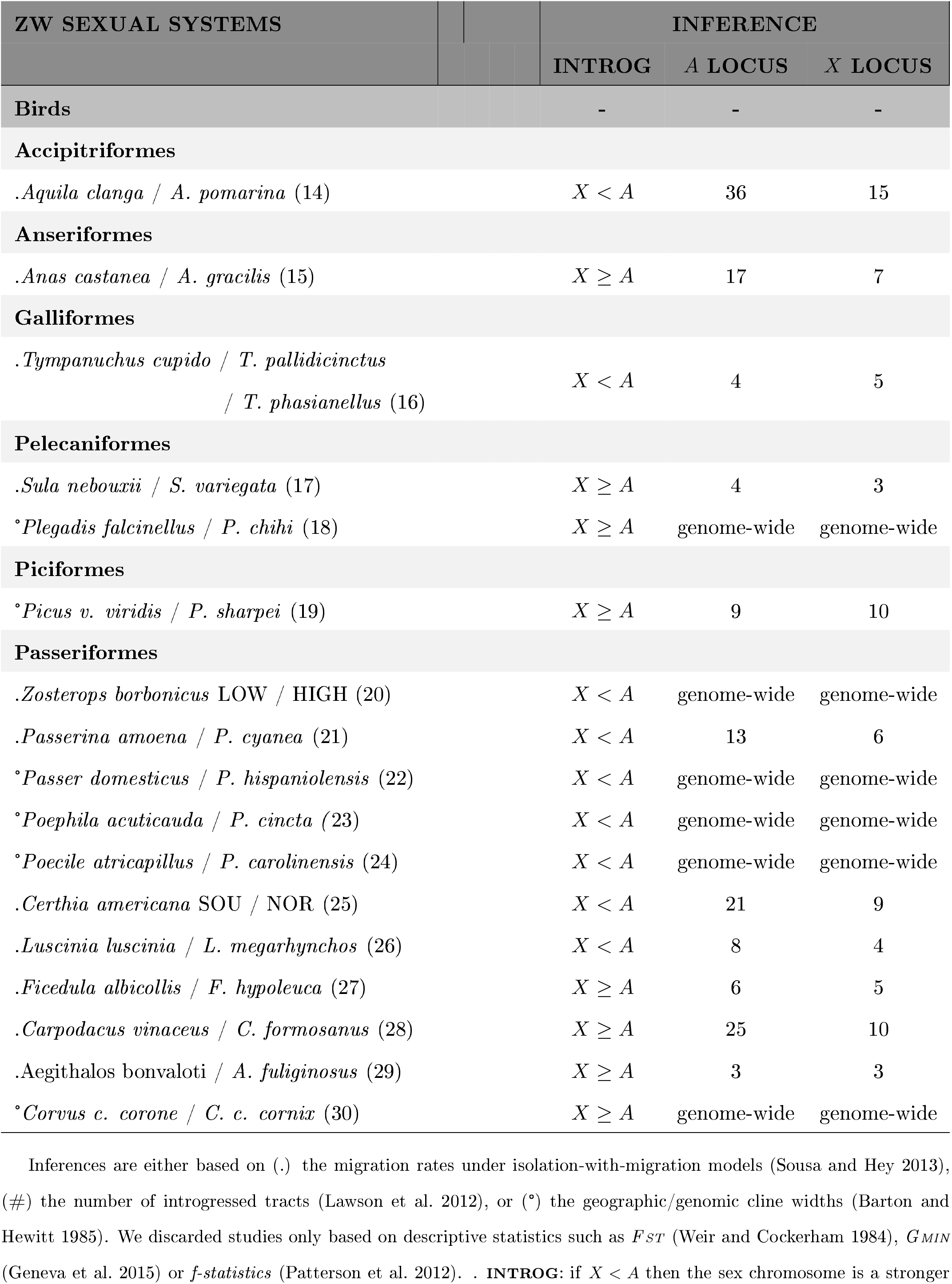
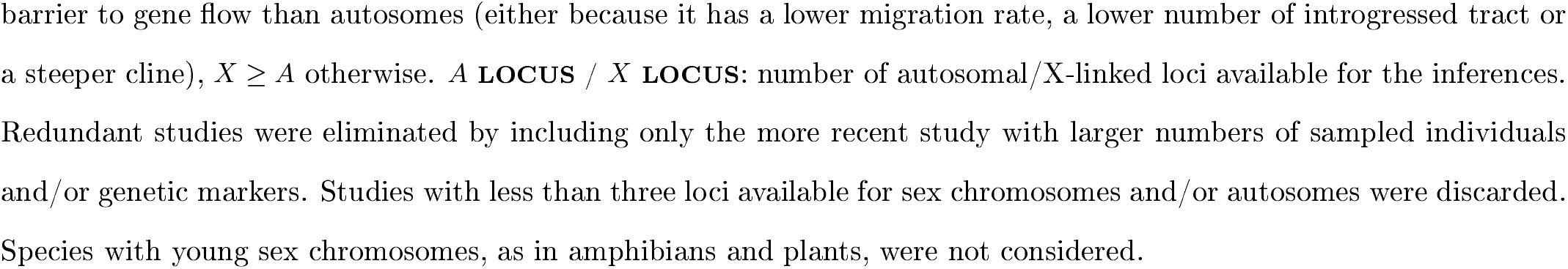
Studies testing whether the sex chromosome is a stronger barrier to gene flow than autosomes.

Overall, patterns are mixed, with ~ 50% of the studies showing lower effective migration rates, steeper clines and/or lower fraction of introgressed tracts on the sex chromosomes compared to the autosomes (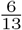 in XY studies and 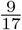 in ZW studies). This apparently inconclusive result hides a strong heterogeneity between clades. In XY systems, almost all mammalian species exhibit stronger X-linked barriers relative to autosomes, in agreement with our theoretical predictions. Dosage compensation by X chromosome inactivation in mammals might contribute to this effect, while the prevalence of male-biased sexual selection would act in the opposite direction. On the contrary, none of the studies in *Drosophila* (except one) shows that the X chromosome is less exchangeable between species than autosomes. This is in line with the fact that the species studied are achiasmatic, although this effect might be counterbalanced by the widespread mechanism of dosage compensation in *Drosophila* (X up-regulation in males). In ZW systems, birds present a balanced mixture of the two patterns across the six orders examined, which prevents us from drawing any definite conclusions. On the one side, the prevalence of male-biased sexual selection in birds is expected to boost the strength of the Z-linked barriers. On the other side, birds may only have partial dosage compensation of Z-linked genes, which would counteracteract this effect. Dosage compensation is highly debated in birds, and more generally, it is unclear how the diverse dosage compensation mechanisms alter the fitness effects of incompatibilities; therefore our interpretations should be regarded as suggestive.

These patterns are preliminary, and should be taken with caution for several reasons. A first caveat is the heterogeneity across studies in both their genetic markers (number and type) and statistical methods. A second concern is the taxonomic sampling, which is overweighted by some clades (e.g. bird species) or some genera within clades (e.g. *Drosophila)*, and likely influenced by positive results; altogether these factors may introduce systematic biases in the global picture. Third, information concerning the mechanisms of dosage compensation or the existence of sex bias is generally reported for a handful of species and then generalized at the level of the order; this precludes a definite test of their impact. Moreover, the balance of power among sex-biased evolutionary forces will depend on their respective magnitudes, which is very hard to measure in nature. Fourth, the effect of sex chromosome linkage, dosage compensation and sex bias on the capacity of sex chromosomes to better resist interspecies introgression may be too subtle to be observed in real data. Actually, the main determinant of the barrier strength between species remains the genetic architecture of reproductive isolation (i.e., the number of barrier loci and their level of recessivity and/or epistasis). Whether or not this differs between sex chromosomes and autosomes remains an open question. Finally, our predictions (which rely on the crucial assumption of rare introgression) may not fully apply to these empirical studies where the frequency of introgressing alleles must be sufficiently high to be detected. Table 3 definitely calls for further comparative analyses that hopefully would provide more robust patterns and a better understanding of their cause.

### Limitations of the model

The present model investigates whether the chromosomal location of reproductive barriers (i.e., sex chromosomes vs. autosomes) influences their capacity to impede interspecies genetic exchange, and ultimately, promote or undermine speciation. However, the rate of accumulation of barrier loci during species divergence may also vary with chromosomal location: the ‘faster-X theory” (Charlesworth et al. 1987) predicts that sex chromosomes (X or Z) evolve more rapidly than autosomes, and as a result, harbor more interspecific barriers. Here, we cannot shed light on the faster-X effect, because the number of reproductive barriers is a parameter of the model; we do not actually model the processes that cause them to accumulate in the first place. Extending our work to model an initial divergence phase would help understand to what extent the faster-X effect contributes to the role of sex chromosomes in speciation.

Our model of genetic exchange in a mainland-island setting rests on the assumption that the rate of migration is low-relative to selection, so that hybrids remain rare. However, to account for the clinal structure of hybrid zones (Barton and Hewitt 1985) and the intermediate frequency of incompatible alleles at their center, it will be necessary to extend our model to more realistic spatial geometries, such as a stepping-stone model or a continuous habitat. This requires different theoretical frameworks that track genomes with multiple introgressed blocks (see Barton 1983, and Barton and Bengtsson 1986), which is beyond the scope of this study. Moreover, in this situation, dominance of the incompatible alleles will become important; thus deviations from the simple expectation that introgression in the sex-linked case is the same as introgression in the autosomal case with appropriately rescaled parameters could arise.

Our analytical treatment is deterministic, i.e., assumes that populations are large enough that genetic drift can be neglected, which is, of course, unrealistic in most natural populations. Also, the outcomes of a hybridization event (where one or a few foreign genomes are introduced and randomly split by recombination) are stochastic (Baird et al. 2003). An extension of the model to include these stochastic processes will thus be necessary to understand the short-term evolution of hybrid genomes.

Another assumption is that all selected alleles are exclusively, and equally, deleterious in the recipient species. A more realistic model would consider allelic effects that vary along the chromosome (i.e., from globally adaptive to locally deleterious). Sachdeva and Barton (2018) studied this scenario by analyzing the introgression dynamics of an autosomal block with linked adaptive and deleterious variants with infinitesimal effects. Importantly, they demonstrated that in this case, deleterious variants can attain high frequency in the recipient species via hitchhiking with genomic blocks with net positive effect (see also Bisschop et al. 2019 for a simulation study). Thus it would be informative to extend the present model to examine how hitchhiking with globally adaptive variants influences the relative barrier strengths of sex chromosomes vs autosomes.

Finally, we would also need to take into account the genome-wide reduction in the effective neutral gene flow due to loosely linked loci (which should affect different chromosomes to similar extents), and would thus further reduce the difference between sex chromosomes and autosomes in species with large genomes.

## DATA AVAILABILITY STATEMENT

SLiM simulation codes for the basic model and notebooks for the maths analyses are available as **Supplementary Codes**:

SuppCode-la: SLiM code for autosomal equilibrium frequencies
SuppCode-lb: SLiM code for X-linked equilibrium frequencies
SuppCode-lc: SLiM code for autosomal barrier strengths (rod configuration)
SuppCode-ld: SLiM code for X-linked barrier strengths (rod configuration)
SuppCode-le: SLiM code for autosomal barrier strengths (embedded configuration)
SuppCode-lf: SLiM code for X-linked barrier strengths (embedded configuration)
SuppCode-2a: Mathematica notebook for equilibrium frequencies
SuppCode-2b: Mathematica notebook for barrier strengths

### Supplementary Texts

1. Appendix A: Distribution of lengths of introgressing blocks
2. Appendix B: Strength of a multilocus barrier to the flow at a neutral marker
3. Model extensions: Epistasis among barrier loci
4. Detailed methods: Individual-based simulations

### Supplementary Figures

Figure SI. Effective number of loci

Figure S2. Correspondence between autosomal and X-linked frequencies

Figure S3. Effect of the number of barrier loci on average equilibrium frequencies and barrier strength at the neutral marker

Figure S4. Effect of very strong coupling on average equilibrium frequencies and barrier strength at the neutral marker Figure S5. Effect of model extensions on average equilibrium frequencies

Figure S6. Effect of model extensions on barrier strength at the neutral marker

Figure S7. Effect of the “rod” vs “embedded” configurations on barrier strength at the neutral marker

### Supplementary Tables

Table SI. Predictions for different evolutionary scenarios

## AUTHOR CONTRIBUTIONS

C.F. designed the project, C.F. and H. S. derived the analytical and numerical results, C.F. conducted the simulations. C.F. and H.S. wrote the manuscript.

## ACKNOWLEDGEMENTS

CF was supported by an Austrian Science Foundation FWF grant (Project M 2463-B29). The computations were performed with the 1ST Austria High-Performance Computing (HPC) Cluster and the Institut Frangais de Bioinformatique (IFB) Core Cluster. We are grateful to Nick Barton and Beatriz Vicoso for critical comments on the model and the manuscript. We also thank Brian Charlesworth, Stuart Baird, and an anonymous reviewer for insightful comments. The authors are aware of no conflicts of interest.

