## Supplemental Information for "The rates of introgression and barriers to genetic exchange between hybridizing species: sex chromosomes vs. autosomes"

### SUPPLEMENTARY TEXT

### 1. Appendix A: Distribution of lengths of introgressing blocks

#### Autosomal introgression

We allow selection coefficients ($s$), migration rates ($m$) and recombination rates ($c$) to be different across the two sexes, thus generalizing the analysis of Barton (1983). In the case of autosomal introgression, the equilibrium proportions of any deleterious block must be exactly the same in males and females in the deterministic limit, even when $s_M \neq s_F$, $m_M \neq m_F$, $c_M \neq c_F$. Let $P(y)$ denote the fraction of diploid individuals (of any sex) carrying a single autosomal deleterious block with $y$ introgressed alleles. At equilibrium, $P(y)$ satisfy the following equations:

$$\begin{aligned} 0 = &\left[\frac{(1-c_M(y-1))e^{-s_M y}}{2} + \frac{(1-c_F(y-1))e^{-s_F y}}{2} - 1\right] P(y) + \sum_{y'=y+1}^{L} (c_M e^{-s_M y'} + c_F e^{-s_F y'}) P(y') \\ &+ [m_F e^{-(s_F/h_F)y} + m_M e^{-(s_M/h_M)y}]\delta_{y,L} \end{aligned} \tag{7}$$

where $\delta_{y,L}$ equals 1 for $y = L$, and 0 otherwise.

Thus $P(y)$ satisfy a set of recursions of the form:

$$0 = -h(y)P(y) + r(y) \sum_{y'=y+1}^{L} k(y')P(y') + f(y)\delta_{y,L} \tag{8}$$

Since recursions of this form also appear in all subsequent analyses presented here, it is useful to provide a general solution. Following Barton (1983) and Barton and Bengtsson (1986), we can re-express these recursions in terms of the function $G(y) = \sum_{y'=y}^{L} k(y)P(y)$, with $P(y) = \frac{G(y)-G(y+1)}{k(y)}$. Then eq. (8) becomes:

$$0 = -\frac{h(y)}{k(y)}(G(y) - G(y+1)) + r(y)G(y+1) \quad \text{for } y < L; \quad G(L) = \frac{f(L)k(L)}{h(L)} \tag{9}$$

These can be solved for $G(y)$ and thence $P(y)$:

$$P(y) = f(L)k(L)r(y) \frac{\prod_{i=1}^{L-1-y} h(i+y) + r(i+y)k(i+y)}{\prod_{i=0}^{L-y} h(i+y)} \quad \text{for } y < L; \quad P(L) = \frac{f(L)}{h(L)} \tag{10}$$

Thus the fraction of diploid individuals (of any sex) carrying a single autosomal deleterious block with $y$ introgressed alleles are:

$$P(y) =(m_F e^{-s_{hom,F}L} + m_M e^{-s_{hom,M}L})(c_M e^{-s_M L} + c_F e^{-s_F L})\times \frac{\prod_{i=1}^{L-1-y} 1 - \frac{(1-c_M(i+y+1))e^{-s_M(i+y)}}{2} - \frac{(1-c_F(i+y+1))e^{-s_F(i+y)}}{2}}{\prod_{i=0}^{L-y} 1 - \frac{(1-c_M(i+y-1))e^{-s_M(i+y)}}{2} - \frac{(1-c_F(i+y-1))e^{-s_F(i+y)}}{2}} \text{ for } y < L \tag{11a}$$

$$P(L) = \frac{m_F e^{-s_{hom,F}L} + m_M e^{-s_{hom,M}L}}{1 - \frac{(1-c_M(L-1))e^{-s_M L}}{2} - \frac{(1-c_F(L-1))e^{-s_F L}}{2}} \tag{11b}$$

where $s_{hom,F} = \frac{s_F}{h_F}$ and $s_{hom,M} = \frac{s_M}{h_M}$.

The frequency of autosomal deleterious blocks is just $P_A(y) = \frac{P(y)}{2}$. When evolutionary processes are slow, i.e., $m_M, m_F, s_M L, s_F L, c_M L, c_F L \ll 1$, such that we can ignore all second and higher order terms in these parameters, this simplifies to:

$$P_A(y) = 2\frac{m_A}{s_A}\theta_A \frac{\prod_{i=1}^{L-y-1} (y+i+1) + \theta_A(y+i)}{\prod_{i=0}^{L-y} (y+i-1) + \theta_A(y+i)} \quad \text{for } y < L; \quad P_A(L) = \frac{m_A}{s_A}\frac{\theta_A}{(L-1)+\theta_A L} \tag{12}$$
$$\text{where} \quad c_A = \frac{c_M + c_F}{2} \quad m_A = \frac{m_M + m_F}{2} \quad s_A = \frac{s_M + s_F}{2}$$

#### X-linked introgression

Let $P_F(y,t)$ and $P_M(y,t)$ denote the fraction of females and males (respectively) carrying a single X-linked deleterious block with $y$ introgressed alleles at time $t$:

$$P_F(y,t+1) = \left(\frac{1-(y-1)c_F}{2}\right) e^{-ys_F} P_F(y,t) + c_F \sum_{y'=y+1}^{L} e^{-y's_F} P_F(y',t) + e^{-ys_M} P_M(y,t) + (m_F e^{-s_{hom,F}y} + m_M e^{-ys_M})\delta_{y,L} \tag{13a}$$

$$P_M(y,t+1) = \left(\frac{1-(y-1)c_F}{2}\right) e^{-ys_F} P_F(y,t) + c_F \sum_{y'=y+1}^{L} e^{-y's_F} P_F(y',t) + m_F e^{-ys_{hom,F}} \delta_{y,L} \tag{13b}$$

where $\delta_{y,L}$ equals 1 for $y = L$, and 0 otherwise; $s_{hom,F} = \frac{s_F}{h_F}$.

Equation (13a) describes how the fraction of females carrying a single X-linked introgressed block with $y$ deleterious alleles changes over time. A female may inherit an entire X-linked block (without any recombination) from her mother (term 1), or inherit a fragment of a larger X-linked block carried by the mother (term 2), or inherit an X-linked block, necessarily without any recombination, from the father (term 3). Equation (13b) for the frequencies of X-linked deleterious blocks in males is similar, except that there is no inheritance from fathers. In addition, for $y = L$, we must account for the fact that the full X-linked block may have been inherited from an immigrant mother or father (final term in eqs. (13a) and (13b)). Since each immigrant female carries two copies of the deleterious block, she must necessarily transmit the entire block to all her offspring (irrespective of whether recombination occurs). Assuming that she has on average $e^{-s_{hom,F}L}$ male and $e^{-s_{hom,F}L}$ female offspring in each generation (where $e^{-s_{hom,F}L}$ is her relative fitness), the frequency of the full X-linked block (with $L$ deleterious alleles) must increase by $m_F e^{-s_{hom,F}L}$ per generation. A similar argument applies for immigrant males, who can transmit the X-linked block only to daughters (at rate $\sim m_M e^{-s_M L}$ per generation).

At equilibrium, we have $P_F(y,t+1) = P_F(y,t) = P_F(y)$ and $P_M(y,t+1) = P_M(y,t) = P_M(y)$. This allows us to solve explicitly for the equilibrium frequencies $P_F(y)$ and $P_M(y)$:

$$P_F(y) = \frac{P_F(y)}{2}(1 - c_F(y-1))e^{-s_F y} + P_M(y)e^{-s_M y} + c_F \sum_{y'=y+1}^{L} P_F(y')e^{-s_F y'} + [m_F e^{-s_{hom,F} y} + m_M e^{-s_M y}]\delta_{y,L} \quad (14a)$$

$$P_M(y) = \frac{P_F(y)}{2}(1 - c_F(y-1))e^{-s_F y} + c_F \sum_{y'=y+1}^{L} P_F(y')e^{-s_F y'} + m_F e^{-s_{hom,F} y}\delta_{y,L} \quad (14b)$$

These can be rewritten as follows:

$$\begin{aligned} 0 = &\left[\frac{e^{-s_F y}(1+e^{-s_M y})(1-c_F(y-1))}{2} - 1\right] P_F(y) + c_F(1+e^{-s_M y}) \sum_{y'=y+1}^{L} P_F(y')e^{-s_F y'} \\ &+ [m_F e^{-s_{hom,F} y}(1+e^{-s_M y}) + m_M e^{-s_M y}]\delta_{y,L} \end{aligned} \quad (15a)$$

$$P_M(y) = \frac{P_F(y) - m_M e^{-s_M y}\delta_{y,L}}{1+e^{-s_M y}} \quad (15b)$$

Equation (15a) is of the general form described above, and can be solved as before to yield the following expressions for $P_F(y)$:

$$\begin{aligned} P_F(y) = &c_F e^{-s_F L}(1+e^{-s_M y})(m_F e^{-s_{hom,F} L}(1+e^{-s_M L}) + m_M e^{-s_M L}) \times \\ &\frac{\prod_{i=1}^{L-1-y} 1 - \frac{e^{-s_F(i+y)}(1+e^{-s_M(i+y)})(1-c_F(y+i+1))}{2}}{\prod_{i=0}^{L-y} 1 - \frac{e^{-s_F(i+y)}(1+e^{-s_M(i+y)})(1-c_F(y+i-1))}{2}} \quad \text{for } y < L \end{aligned} \quad (16a)$$

$$P_F(L) = \frac{m_F e^{-s_{hom,F} L}(1+e^{-s_M L}) + m_M e^{-s_M L}}{1 - \frac{e^{-s_F L}(1+e^{-s_M L})(1-c_F(L-1))}{2}} \quad (16b)$$

The corresponding block frequencies in males $P_M(y)$ and the average frequency of X-linked blocks in the population

$P_X(y) = \frac{P_M(y)+P_F(y)}{3}$ can be obtained from these using eq. (15b).

**Continuous time limit:**

In the limit where all evolutionary processes are slow (i.e., $m_M, m_F, s_M L, s_F L, c_F L \ll 1$), such that genotype frequencies change very little in a single generation, it is useful to further approximate equations (13) by the corresponding continuous time equations. Formally, this can be done by substituting: $m \to m\delta t$, $s \to s\delta t$, $c \to c\delta t$ and so on, where $\delta t$ is an infinitesimal time interval. Then by Taylor expanding equations (13) in powers of $\delta t$ and retaining only lowest order terms in $\delta t$, we obtain:

$$\frac{\partial P_F(y)}{\partial t} = -\left[\frac{2s_F}{3}y + \frac{s_M}{3}y + \frac{2c_F}{3}(y-1)\right] P_F(y) + \frac{4c_F}{3}\sum_{y'=y+1}^{L} P_F(y') + (\frac{4m_F}{3} + \frac{2m_M}{3})\delta_{y,L} \tag{17a}$$

$$P_M(y,t) \sim \frac{P_F(y,t)}{2} \tag{17b}$$

The fact that $P_M(y,t) \sim \frac{P_F(y,t)}{2}$ implies that the total frequency of any X-linked block, given by $P_X(y,t) = \frac{P_F(y,t)+P_M(y,t)}{3}$ is approximately $\frac{P_F(y,t)}{2}$. Thus, we have:

$$\begin{aligned}\frac{\partial P_X(y)}{\partial t} &= -\left[s_X y + c_X(y-1)\right] P_X(y) + 2c_X \sum_{y'=y+1}^{L} P_X(y') + m_X \delta_{y,L} \\ \text{where} \quad c_X &= \frac{2c_F}{3} \quad m_X = \frac{2m_F + m_M}{3} \quad s_X = \frac{2s_F + s_M}{3}\end{aligned} \tag{18}$$

Equation (18) is identical in form to the corresponding equation for the dynamics of the block length distribution in the autosomal case (Barton 1983) (see also eq. (1) in the main text), but with $s_A$, $m_A$ and $c_A$ replaced by the sex-averaged selection coefficient $s_X$, sex-averaged migration rate $m_X$ and sex-averaged recombination rate $c_X$ (which are weighted sums of male and female contributions). Equation (18) can be solved to obtain an exact time-dependent solution for $P_X(y,t)$ (see Appendix in Baird 1995).

However, it is more useful to consider equilibrium block frequencies $P_X(y)$, obtained by setting the time derivative in equation (18) to zero. The resultant equations can be solved recursively, as in Barton (1983), by first re-expressing it in terms of the cumulative distribution $G_X(y) = \sum_{y'=y}^{L} P_X(y')$, then solving recursively for $G_X(y)$, and finally using this solution to obtain $P_X(y) = G_X(y) - G_X(y+1)$. This yields the following equilibrium proportions of different X-linked blocks:

$$P_X(y) = 2\frac{m_X}{s_X}\theta_X \frac{\prod_{i=1}^{L-y-1}(y+i+1)+\theta_X(y+i)}{\prod_{i=0}^{L-y}(y+i-1)+\theta_X(y+i)} \quad \text{for } y < L; \quad P_X(L) = \frac{m_X}{s_X}\frac{\theta_X}{(L-1)+\theta_X L} \tag{19}$$

where $\theta_X = \frac{s_X}{c_X}$ is a measure of the strength of coupling between deleterious alleles on the X chromosome. This can be used to calculate the average frequency of deleterious alleles, averaged over all loci on the X chromosome: $\overline{p}_X = \sum_{y=1}^{L} P_X(y)\frac{y}{L}$.

Note that in writing eq. (13), we assumed that deleterious alleles have arbitrary dominance coefficient $h$ (such that females who are homozygous vs. heterozygous for the full deleterious block have fitness $e^{-s_{hom,F}L}$ and $e^{-s_F L}$ respectively, where $s_{hom,F} = \frac{s_F}{h_F}$). However, retaining only lowest order terms in $s_F$, $s_M$, $c_F$, $m_F$ and $m_M$ results in the approximations: $m_F e^{-s_{hom,F}L} \sim m_F$, and $m_M e^{-s_M L} \sim m_M$. Thus, the final equation (18) is independent of the dominance coefficient of deleterious alleles, as long as net selection against immigrant genotypes, $S = sL$, is sufficiently weak. For larger values of $S$, we expect $P_F(y)$ and $P_M(y)$ to depend on $h$, as can be seen from the full solution in eq. (16) above.

### 2. Appendix B: Strength of a multilocus barrier to the flow at a neutral marker

#### Autosomal barrier

We consider the rod model where the neutral marker is on one side of the deleterious block. Then assuming that deleterious blocks remain rare in the population, the neutral marker will always be found within one of $L + 1$ possible genetic backgrounds, where each such background is defined by the number $y$ of deleterious alleles that it contains. Moreover, in the case of autosomal introgression, the equilibrium proportions of any genetic background must be exactly the same in males and females in the deterministic limit, irrespective of whether parameters are unequal across sexes. Let $U(y)$ denote the fraction of individuals (of any sex) carrying a single introgressed block which contains the neutral marker in conjunction with $y$ deleterious alleles. At equilibrium, we have:

$$0 = \left[\frac{(1-c_M(y-1+\alpha))e^{-s_M y}}{2} + \frac{(1-c_F(y-1+\alpha))e^{-s_F y}}{2} - 1\right] U(y) + \sum_{y'=y+1}^{L} \left(\frac{c_M}{2}e^{-s_M y'} + \frac{c_F}{2}e^{-s_F y'}\right)U(y') \\ + [m_F e^{-s_{hom,F}y} + m_M e^{-s_{hom,M}y}]\delta_{y,L} \tag{20}$$

where $\delta_{y,L}$ equals 1 for $y = L$, and 0 otherwise; $s_{hom,F} = \frac{s_F}{h_F}$ and $s_{hom,M} = \frac{s_M}{h_M}$.

The barrier strength can be found as:

$$b = \frac{m_F + m_M}{\delta U_0} \quad \text{where:} \quad \delta U_0 = \alpha \sum_{y'=1}^{L} \left(\frac{c_M}{2}e^{-s_M y'} + \frac{c_F}{2}e^{-s_F y'}\right)U(y') \tag{21}$$

The recursions in (20) have the form described above, and can be solved to determine $U(y)$ for all $y$. Substituting these expressions for $U(y)$ into eq. (21) yields the following expression for barrier strength:

$$b = \frac{1 + \frac{m_M}{m_F}}{e^{-s_{hom,F}L} + \frac{m_M}{m_F}e^{-s_{hom,M}L}} \left(\frac{2-(1-c_M(L-1+\alpha))e^{-s_M L} - (1-c_F(L-1+\alpha))e^{-s_F L}}{\alpha[c_M e^{-s_M L} + c_F e^{-s_F L}]}\right) \times \\ \prod_{y=1}^{L-1} \left[\frac{2-(1-c_M(y-1+\alpha))e^{-s_M y} - (1-c_F(y-1+\alpha))e^{-s_F y}}{2-(1-c_M(y+\alpha))e^{-s_M y} - (1-c_F(y+\alpha))e^{-s_F y}}\right] \tag{22}$$

Note that the barrier strength is independent of the strength of migration (as it should be), and depends only on the asymmetry in migration rates between the two sexes. In the absence of sex-specificities, i.e., for $s_M = s_F = s$, $m_M = m_F = m$, $c_M = c_F = c$ and $h_M = h_F = h$, the expression for barrier strength reduces to a simpler form:

$$b = e^{s_{hom} L} \left( \frac{1 - (1 - c(L + \alpha))e^{-sL}}{\alpha c e^{-sL}} \right) \prod_{y=1}^{L} \left[ \frac{1 - (1 - c(y - 1 + \alpha))e^{-sy}}{1 - (1 - c(y + \alpha))e^{-sy}} \right] \tag{23}$$

This is identical to the expression in eq. $A3$ of Barton and Bengtsson (1986) except for the extra term $e^{s_{hom} L}$, which accounts for the fact that immigrants suffer an initial (i.e., first-generation) disadvantage which is proportional to $e^{-s_{hom} L}$ (or $e^{-sL}$ in the case of the haploid model considered by Barton and Bengtsson 1986).

In the limit where all evolutionary processes are slow, such that second order terms in $m$, $s$ and $c$ can be neglected, the expression in eq. (22) simplifies to:

$$b_A = \frac{\Gamma\left(L + \frac{\alpha + \theta_A}{1 + \theta_A}\right) \Gamma\left(\frac{\alpha}{1 + \theta_A}\right)}{\Gamma\left(L + \frac{\alpha}{1 + \theta_A}\right) \Gamma\left(\frac{\alpha + \theta_A}{1 + \theta_A}\right)} \tag{24}$$

where $\theta_A = \frac{s_A}{c_A}$ is a measure of the strength of coupling between locally deleterious alleles on the autosomes, and $\alpha$ is a measure of the closeness of the neutral marker to the selected block compared to the genetic length of the latter.

#### X-linked barrier

Let $U_F(y, t)$ and $U_M(y, t)$ denote the fraction of females and males carrying an X-linked introgressed block which contains the neutral marker in conjunction with $y$ deleterious alleles at time $t$:

$$\begin{aligned} U_F(y, t+1) = &\left( \frac{1 - c_F(\alpha + y - 1)}{2} \right) e^{-y s_F} U_F(y, t) + \frac{c_F}{2} \sum_{y'=y+1}^{L} e^{-y' s_F} U_F(y', t) + e^{-y s_M} U_M(y, t) + \\ &(m_F e^{-s_{hom,F} y} + m_M e^{-y s_M}) \delta_{y,L} + \left( \frac{\alpha c_F}{2} \sum_{y'=1}^{L} e^{-y' s_F} U_F(y', t) \right) \psi_{y,0} \end{aligned} \tag{25a}$$

$$\begin{aligned} U_M(y, t+1) = &\left( \frac{1 - c_F(\alpha + y - 1)}{2} \right) e^{-y s_F} U_F(y, t) + \frac{c_F}{2} \sum_{y'=y+1}^{L} e^{-y' s_F} U_F(y', t) + m_F e^{-s_{hom,F} y} \delta_{y,L} \\ &+ \left( \frac{\alpha c_F}{2} \sum_{y'=1}^{L} e^{-y' s_F} U_F(y', t) \right) \psi_{y,0} \end{aligned} \tag{25b}$$

where $\delta_{y,L}$ equals 1 for $y = L$, and 0 otherwise; $\psi_{y,0}$ equals 1 for $y = 0$, and 0 otherwise; $s_{hom,F} = \frac{s_F}{h_F}$.

Equation (25a) describes how the fraction of females carrying a single X-linked neutral marker associated to an introgressed block with $y$ deleterious alleles changes over time; and equation (25b) is the equivalent recursion for males. These equations are of the same form as (13a) and (13b), except that we must account for the fact that the neutral marker can be associated with the recipient background, i.e. $y = 0$ (final term in eqs. (25a) and (25b)).

At equilibrium, we have $U_F(y, t+1) = U_F(y, t) = U_F(y)$ and $U_M(y, t+1) = U_M(y, t) = U_M(y)$. This allows us to solve explicitly for the equilibrium frequencies $U_F(y)$ and $U_M(y)$:

$$U_F(y) = \frac{U_F(y)}{2}(1 - c_F(y-1+\alpha))e^{-s_F y} + U_M(y)e^{-s_M y} + \frac{c_F}{2}\sum_{y'=y+1}^{L} U_F(y')e^{-s_F y'} + [m_F e^{-s_{hom,F} y} + m_M e^{-s_M y}]\delta_{y,L} \tag{26a}$$

$$U_M(y) = \frac{U_F(y)}{2}(1 - c_F(y-1+\alpha))e^{-s_F y} + \frac{c_F}{2}\sum_{y'=y+1}^{L} U_F(y')e^{-s_F y'} + [m_F e^{-s_{hom,F} y}]\delta_{y,L} \tag{26b}$$

These can be rewritten as:

$$\begin{aligned} 0 = &\left[\frac{e^{-s_F y}(1+e^{-s_M y})(1-c_F(y-1+\alpha))}{2} - 1\right] U_F(y) + \frac{c_F}{2}(1+e^{-s_M y}) \sum_{y'=y+1}^{L} U_F(y')e^{-s_F y'} \\ &+ [m_F e^{-s_{hom,F} y}(1+e^{-s_M y}) + m_M e^{-s_M y}]\delta_{y,L} \end{aligned} \tag{27}$$

$$U_M(y) = \frac{U_F(y) - m_M e^{-s_M y}\delta_{y,L}}{1+e^{-s_M y}} \tag{28}$$

In this case, the barrier strength is given by:

$$b_X = \frac{m_F + \frac{m_M}{2}}{\frac{\delta U_{0,F} + \delta U_{0,M}}{2}} \quad \text{where} \quad \delta U_{0,F} = \delta U_{0,M} = \frac{\alpha}{2} c_F \sum_{y=1}^{L} U_F(y) e^{-s_F y} \tag{29}$$

This can be calculated explicitly by first solving eq. (26a) for $U_F(y)$ and then substituting into the expression for $b_X$. This yields the following expression for barrier strength:

$$\begin{aligned} b_X = &\frac{1 + \frac{m_M}{2m_F}}{e^{-s_{hom,F} L}\frac{(1+e^{-s_M L})}{2} + \frac{m_M}{2m_F}e^{-s_M L}} \left(\frac{2 - e^{-s_F L}(1+e^{-s_M L})(1-c_F(L-1+\alpha))}{2\alpha c_F e^{-s_F L}}\right) \times \\ &\prod_{y=1}^{L-1}\left[\frac{2 - e^{-s_F y}(1+e^{-s_M y})(1-c_F(y-1+\alpha))}{2 - e^{-s_F y}(1+e^{-s_M y})(1-c_F(y+\alpha))}\right] \end{aligned} \tag{30}$$

As before, to lowest order in $s$, $m$ and $c$, this reduces to a much simpler expression:

$$b_X = \frac{\Gamma\left(L + \frac{\alpha+\theta_X}{1+\theta_X}\right)\Gamma\left(\frac{\alpha}{1+\theta_X}\right)}{\Gamma\left(L + \frac{\alpha}{1+\theta_X}\right)\Gamma\left(\frac{\alpha+\theta_X}{1+\theta_X}\right)} \tag{31}$$

where $\theta_X = \frac{s_X}{c_X}$ is a measure of the strength of coupling between locally deleterious alleles on the X chromosome, and $\alpha$ is a measure of the closeness of the neutral marker to the selected block compared to the genetic length of the latter.

**Connection with the results of Muirhead and Presgraves (2016):**

Here we demonstrate that in the limit of very tight linkage between barrier loci, our results for barrier strength reduce to those of Muirhead and Presgraves (2016), who consider a neutral marker linked to a single incompatible allele. To make this connection, it is useful to rewrite eq. (31) in terms of $r = \alpha c_F = (2/3)\alpha c_X$, where $r$ is the rate of recombination between the neutral marker and the nearest selected locus on the $X$ in females:

$$b_X = \frac{\Gamma\left[L + \frac{(2/3)(r/c_X)+(s_X/c_X)}{1+(s_X/c_X)}\right]\Gamma\left[\frac{(2/3)(r/c_X)}{1+(s_X/c_X)}\right]}{\Gamma\left[L + \frac{(2/3)(r/c_X)}{1+(s_X/c_X)}\right]\Gamma\left[\frac{(2/3)(r/c_X)+(s_X/c_X)}{1+(s_X/c_X)}\right]} \tag{32}$$

In the limit of very tight linkage between loci, i.e., $c_X \to 0$, the above expression reduces to $b_X \sim 1 + (3/2)(S_X/r)$, where $S_X = s_X L$.

Muirhead and Presgraves (2016) calculate $p_{ef}$ and $p_{em}$, the permeabilities (i.e., the inverse of the barrier strength) for a neutral allele linked to a single incompatible allele introduced by a female migrant and a male migrant respectively (see their equations 3 and 4). These can be used to calculate the total barrier strength as follows: $b_X = \frac{2}{3}\frac{1}{p_{ef}} + \frac{1}{3}\frac{1}{p_{em}}$. In the limit where evolutionary forces are weak, i.e., $s, r \ll 1$, such that it is sufficient to retain only lowest order terms in $s$ and $r$ in their expressions for $p_{ef}$ and $p_{em}$, the X-linked barrier strength again becomes $b_X \sim 1 + (3/2)(s_X/r)$. Here $s_X$ is the sex-averaged X-linked selective effect which, in their notation, is simply: $s_X = \frac{2}{3}hs + \frac{1}{3}s$.

### 3. Model extensions: Epistasis among barrier loci

We have assumed in the main text that selection acts independently (i.e., multiplicatively) against incompatible introgressing alleles. However, deleterious interaction between them (i.e. negative epistasis, $\varepsilon < 1$) is an important component of speciation, and is the basis of the seminal DMI model (Dobzhansky 1937; Muller 1940). Therefore, we now consider the case where a fraction $\beta$ of the loci act epistatically, and a fraction $(1-\beta)$ acts multiplicatively. Following Barton and Bengtsson (1986), the fitness of an individual with $y$ introgressed alleles is then $e^{-ay-by^2}$, where $a = (1-\beta+\beta\varepsilon)s$, $b = \beta s(1-\varepsilon)$ and $\varepsilon$ gives the direction and strength of pairwise epistasis (note that when $\beta = 0$, the fitness of an individual carrying a block with $y$ equal-effect loci is $v(y) = e^{-ys}$). When $\beta > 0$ and $\varepsilon < 1$, the fitness of an incompatible block decreases non-linearly with the number of barrier loci it carries (note that this is quite different from the kind of epistasis considered in the DMI model).

We evaluated a version of the model where all loci act epistatically ($\beta = 1$), and found that epistasis acts by strongly decreasing the equilibrium frequency of both autosomes and sex chromosomes as a function of the number of incompatible loci (when $L \geq 1$, blue and red; Figure S5D). Accordingly, the strength of an epistatic multilocus barrier is much stronger than in the multiplicative case (by a factor $\sim 10^4$ with $L = 100$, $\alpha = 0.01$ and $\varepsilon = 0.1$; Figure S6D). Importantly, and contrary to the multiplicative model, there is a deficit of autosomal to sex-linked introgression when the number of deleterious loci is sufficiently large (say $L \gg 10$), which amplifies with the number of incompatible alleles (blue and red; Figure S5D). The ratio of the equilibrium frequencies between the two chromosome types can be as large as $\frac{\overline{p}_X}{\overline{p}_A} \sim 12$ (with $L = 100$ and $\varepsilon = 0.1$, red; Figure S5D), which leads to an autosomal barrier strength $\sim 30$ times higher than that on the sex chromosome (with $L = 100$, $\alpha = 0.01$ and $\varepsilon = 0.1$, red; Figure S6D). This happens because with epistasis, the decreased fitness of immigrants and F1 hybrids, which carry entire incompatible blocks, prevails over that of later generation hybrids (which carry shorter incompatible blocks); and considering that immigrant alleles are selected against more heavily on autosomes relative to sex chromosomes due to the weaker (hemizygous) selection acting on migrant XY males (in the absence of dosage compensation).

### 4. Detailed methods: Individual-based simulations

Simulations were performed with SLiM 3.3 (Haller and Messer 2019) following a Wright–Fisher life cycle. Two constant-size populations of $N = 100,000$ sexually-reproducing diploid individuals were simulated with discrete and non-overlapping generations. For simplicity, we assume a male-heterogametic sexual system (males are XY, and females are XX) with $1:1$ sex-ratio. We model steady migration from the donor into the recipient species by replacing a fraction $m_F$ of females and a fraction $m_M$ of males by immigrants in each generation. Offspring are then generated by drawing a random mother and a random father according to their fitness. The offspring's genotype is created from the chosen parents as follows: the first offspring haploid genome is produced via recombination between the two genomes of the female, and the second via recombination between the two genomes of the male (except in the case of the X which do not recombine in males). Once all offspring are generated, the offspring generation becomes the new parental generation. Note that, unlike in the analytical treatment, an individual might bear multiple introgressed fragments.

Simulated genomic blocks, lying either on an autosome or a X chromosome, carry $L$ equally spaced loci, which are fixed for different alleles in the two species at the start of the simulations. Recombination occurs at a uniform rate $c_F$ per locus per generation in females ($c_M$ in males). The expected number of crossovers per generation, within the entire block, is thus $C = c(L-1)$. For each offspring genome, the number of recombination breakpoints is drawn from a Poisson distribution with mean equal to $c(L-1)$ and their positions are chosen by sampling uniformly along the block. We assume null recombination between the X and the Y chromosomes in males. Note that the simulation model can be extended to include epistasis by changing the form of the fitness function.

We followed the average frequency of the deleterious alleles in the recipient species across generations, separately for sex chromosomes (i.e. $\overline{p}_X$) and autosomes (i.e. $\overline{p}_A$). Simulations were run for $t = 10,000$ generations, which is more than sufficient for migration-selection-recombination balance to be reached. For a given set of parameters, we performed 100 replicate simulations. We then investigate the effect of the selected block as a barrier against neutral gene-flow. Once migration-selection balance is reached in the recipient species (in practice, after $t = 10,000$ generations), we introduced a neutral marker differentially fixed between them at the extremity ("rod" configuration) or in the center ("embedded" configuration) of the $L$-locus block. The recombination rate of the neutral marker with the nearest deleterious variant (or with each of the two surrounding deleterious variants) on the selected block is $\alpha c$. We calculated the effective migration rate of the neutral allele at equilibrium by monitoring its rate of increase in the recipient species: $m_e = \frac{\overline{u}_{recipient}(t+1)-\overline{u}_{recipient}(t)}{\overline{u}_{donor}-\overline{u}_{recipient}(t)}$, where $\overline{u}_{recipient}(t)$ and $\overline{u}_{donor}$ are the average frequencies of the neutral marker in the recipient and donor species, respectively (see Figure S3A). Note that $\overline{u}_{donor} = 1$, since the donor species is fixed for the neutral allele, and migration is one-way. In practice, we fitted a linear regression to the log of $\overline{u}_{recipient}(t)$ as a function of the number of generations to estimate its slope (i.e. $m_e$) using the R function "lm" (from package "stats" v3.6.1, R 2013). We calculated its confidence interval using the R function "confint" from the same package. We based our calculation on the first few hundred generations that showed a constant rate of increase of the neutral allele.

### SUPPLEMENTARY FIGURES

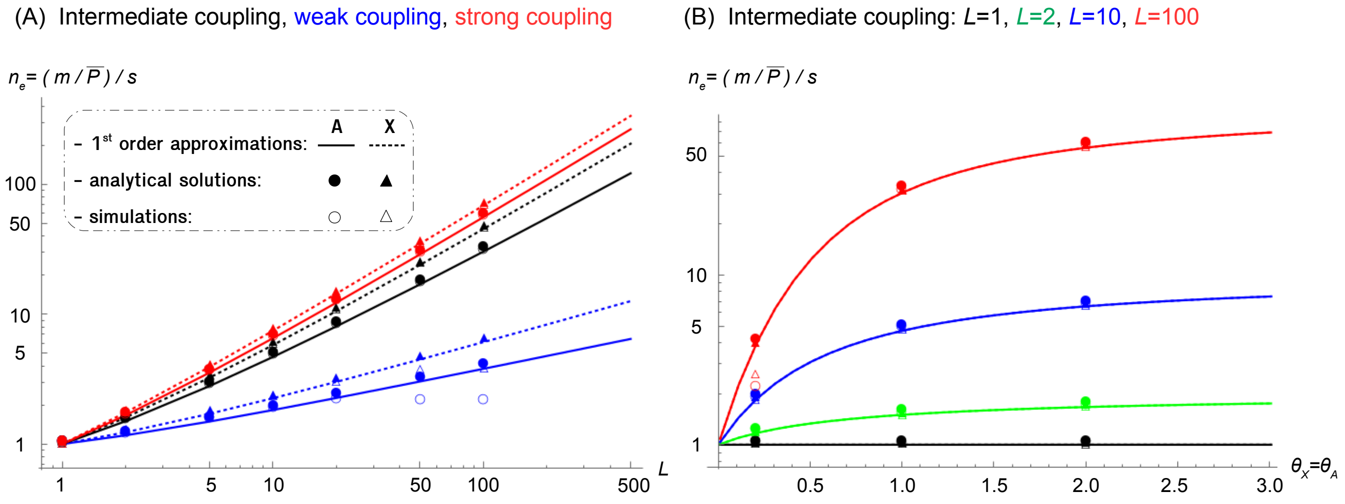


**Figure S1. Effective number of loci**

**(A)** Effective number of loci, $n_e$, plotted against the number of selected loci on the genomic block, $L$. Colors stand for values of the coupling coefficient: $\theta$ small ($cL = 0.25$, $\theta_X = 0.3$, $\theta_A = 0.2$; blue), $\theta$ intermediate ($cL = 0.05$, $\theta_X = 1.5$, $\theta_A = 1$; black) and $\theta$ large ($cL = 0.025$, $\theta_X = 3$, $\theta_A = 2$; red). **(B)** Effective number of loci, $n_e$, plotted against the coupling coefficient, $\theta$. Parameters are scaled such that $\theta_X = \theta_A$ and $\frac{m_X}{S_X} = \frac{m_A}{S_A}$, which leads to $n_{eX} = n_{eA}$. Colors stand for the number of selected loci on the block: $L = 1$ (black), $L = 2$ (green), $L = 10$ (blue) and $L = 100$ (red). **A**utosomes are depicted with solid lines, ● and ○; **X** chromosomes with dotted lines, ▲ and △. Parameter values are: $h = 0.5$, $m = 0.001$, $sL = 0.05$, $m_X/S_X = m_A/S_A = 0.02$ and $N = 10^5$ (simulations).

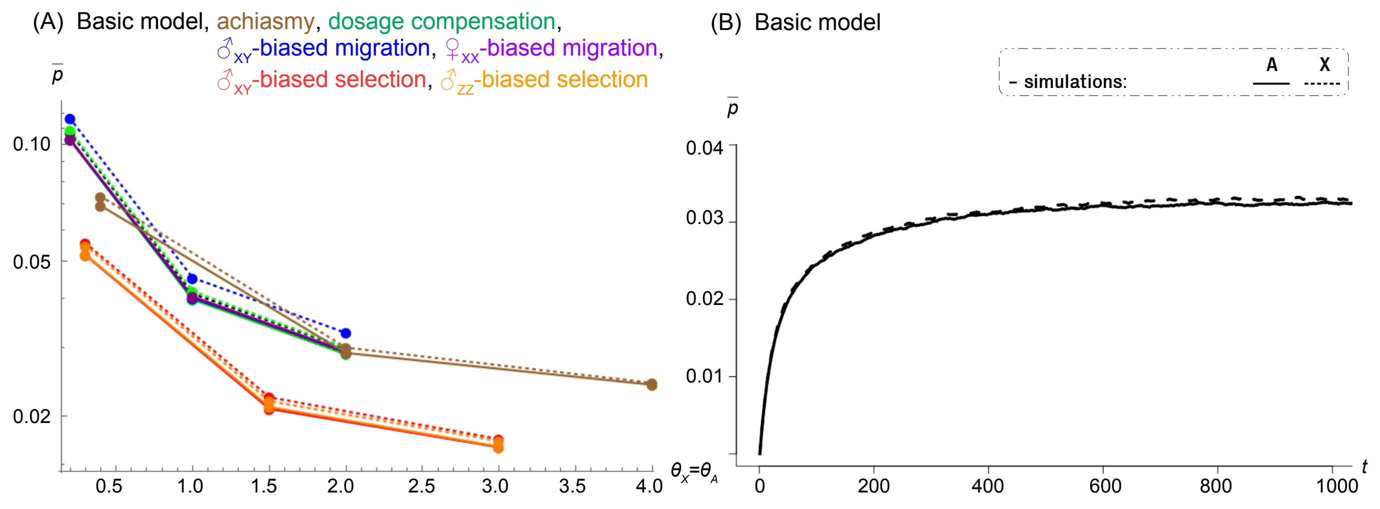


**Figure S2. Correspondence between autosomal and X-linked frequencies**

Average frequency of the deleterious alleles in the recipient species (**A**utosomal, solid lines: $\overline{p}_A$; **X**-linked, dotted lines: $\overline{p}_X$) plotted against: **(A)** the coupling coefficient, $\theta$; **(B)** time in generations, $t$. Parameters are scaled such that $\theta_X = \theta_A$ and $\frac{m_X}{S_X} = \frac{m_A}{S_A}$, which leads to $\overline{p}_X = \overline{p}_A$ in the equilibrium distribution (A), and to $\overline{p}_X(t) = \overline{p}_A(t)$ in the dynamics (B). Colors stand for a range of different scenarios: basic model (in (A): $\theta_X = \theta_A = \{0.2, 1, 2\}$, $m_X/S_X = m_A/S_A = 0.02$; in (B): $\theta_X = \theta_A = 1.5$, $m_X/S_X = m_A/S_A = 0.02$; black), dosage compensation ($\theta_X = \theta_A = \{0.2, 1, 2\}$, $m_X/S_X = m_A/S_A = 0.02$; green), XY male-biased migration ($\theta_X = \theta_A = \{0.2, 1, 2\}$, $m_X/S_X = m_A/S_A = 0.02$; blue), XX female-biased migration ($\theta_X = \theta_A = \{0.2, 1, 2\}$, $m_X/S_X = m_A/S_A = 0.02$; purple), XY male-biased selection ($\theta_X = \theta_A = \{0.3, 1.5, 3\}$, $m_X/S_X = m_A/S_A = 0.013$; red), ZZ male-biased selection ($\theta_X = \theta_A = \{0.3, 1.5, 3\}$, $m_X/S_X = m_A/S_A = 0.013$; orange) and achiasmy ($\theta_X = \theta_A = \{0.4, 2, 4\}$, $m_X/S_X = m_A/S_A = 0.02$; brown). Note that the number of selected loci on the block is fixed to $L = 10$, $h = 0.5$, $N = 10^5$ and that only individual-based simulations are shown. Simulations were run for (A) 10,000 generations; (B) 1,000 generations.

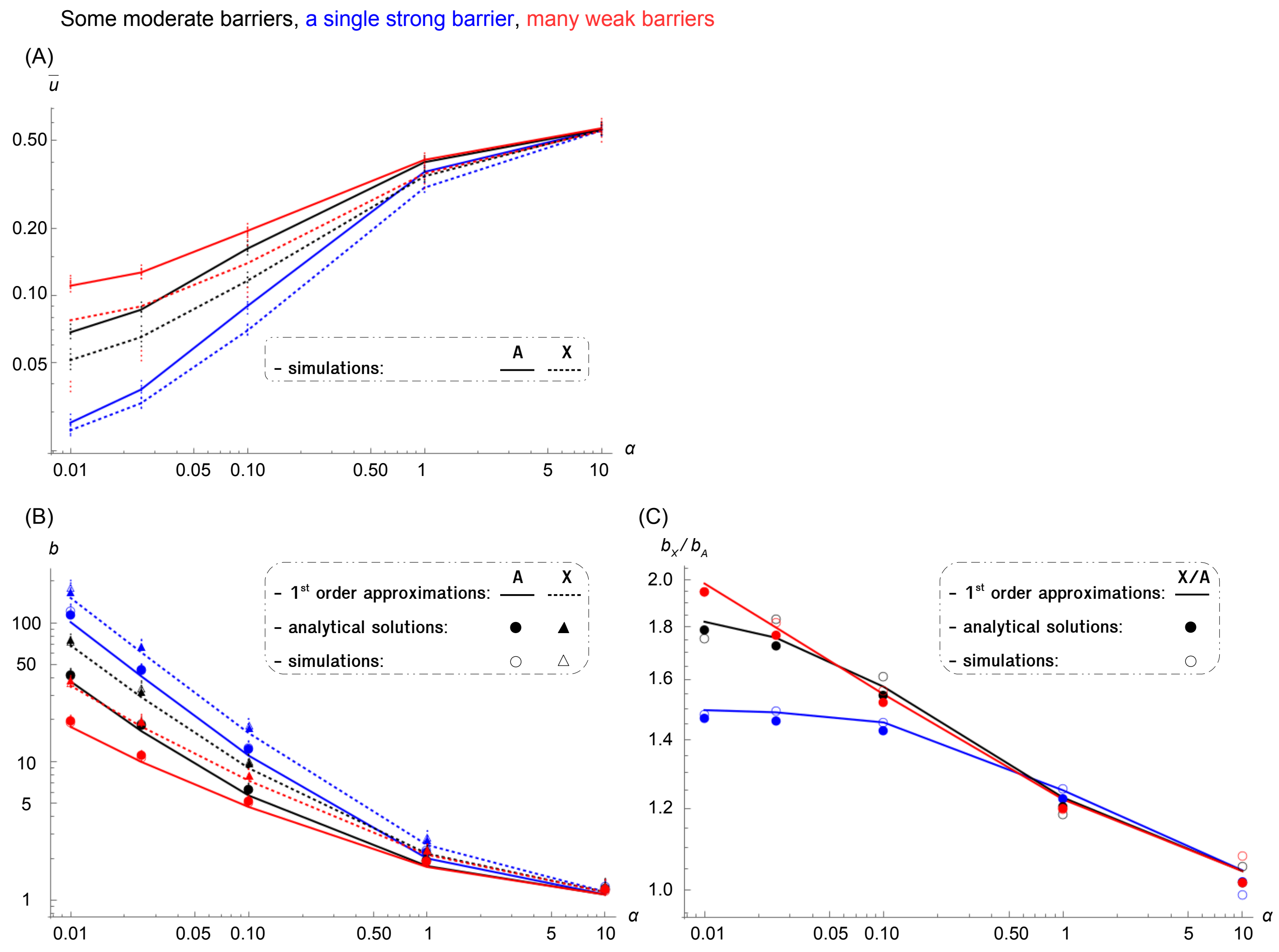


**Figure S3. Effect of the number of barrier loci on average equilibrium frequencies and barrier strength at the neutral marker**

Quantities are plotted against the proximity of the neutral marker to its nearest selected locus, $\alpha$. **(A)** Simulated average frequency of the neutral allele in the recipient species after 10,000 generations (**A**utosomal: $\overline{u}_A$; **X**-linked: $\overline{u}_X$). **(B)** Barrier strength at the neutral marker (**A**utosomal: $b_A$; **X**-linked: $b_X$). **(C)** the $\frac{b_X}{b_A}$ ratio. Colors stand for values of the number of selected loci on the genomic block, $L$: a single strongly selected locus (blue), ten moderately selected loci (black) and one hundred weakly selected loci (red). Parameter values are: $h = 0.5$, $m = 0.001$, $cL = 0.05$, $sL = 0.05$, $m_X/S_X = m_A/S_A = 0.02$, $\theta_X = 1.5$, $\theta_A = 1$ and $N = 10^5$ (simulations). Other details match Figure 2.

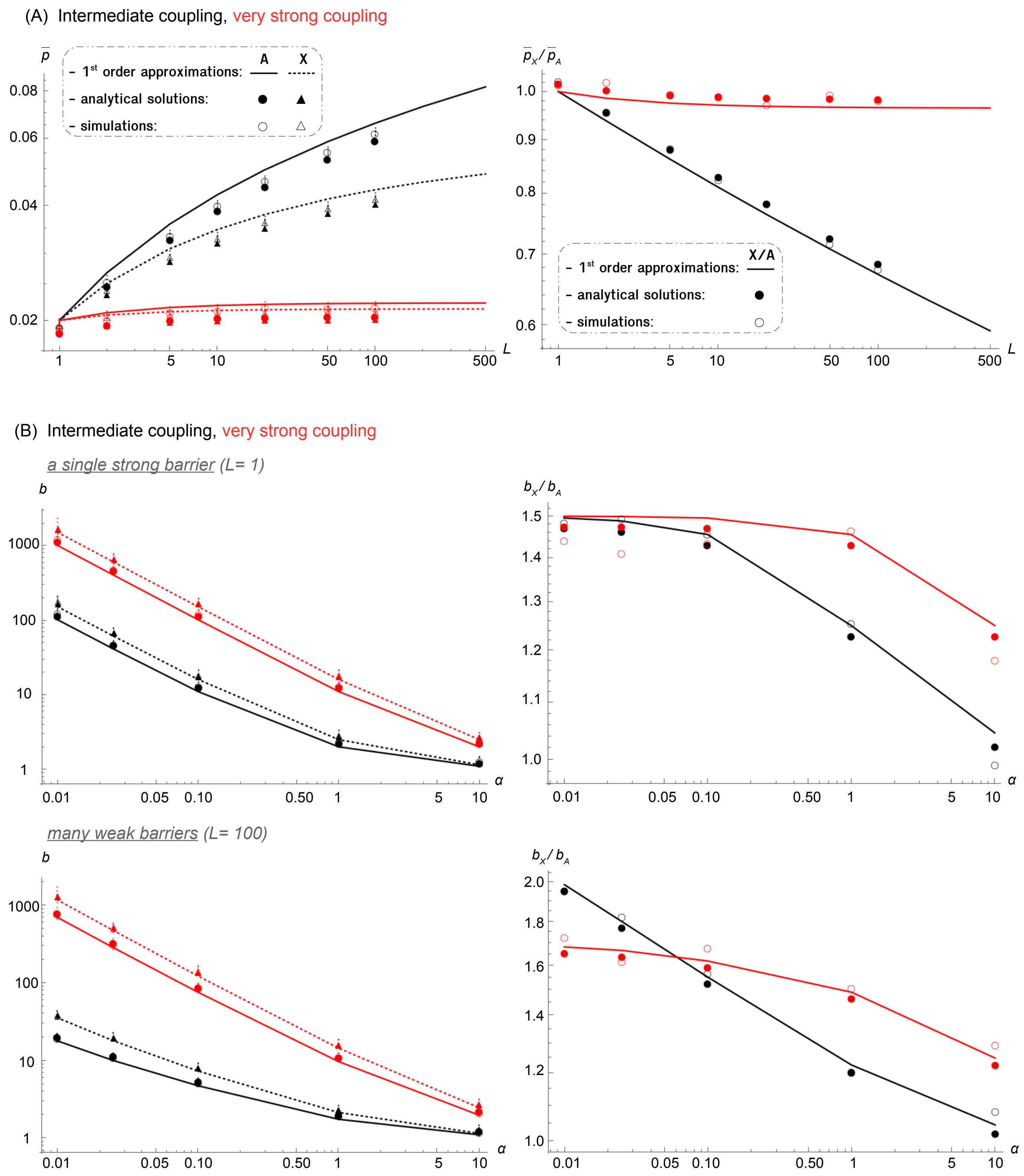


**Figure S4. Effect of very strong coupling on average equilibrium frequencies and barrier strength at the neutral marker**

**(A)** Average frequency of the deleterious alleles in the recipient species (**A**utosomal: $\overline{p}_A$; **X**-linked: $\overline{p}_X$; their ratio: $\frac{\overline{p}_X}{\overline{p}_A}$) plotted against the number of selected loci on the genomic block, $L$. **(B)** Barrier strength at the neutral marker

(**A**utosomal: $b_A$; **X**-linked: $b_X$; their ratio: $\frac{b_X}{b_A}$) plotted against its proximity to the nearest selected locus, $\alpha$. Colors stand for values of the composite parameter $\theta = \frac{s}{c}$ at the selected loci. Parameter values for intermediate $\theta$ are: $h = 0.5$, $m = 0.001$, $cL = 0.05$, $sL = 0.05$, $m_X/S_X = m_A/S_A = 0.02$, $\theta_X = 1.5$, $\theta_A = 1$ and $N = 10^5$ (simulations). Parameters for very strong coupling: $cL = 0.005$, $\theta_X = 15$, $\theta_A = 10$. Other details match Figures 1 and 2.

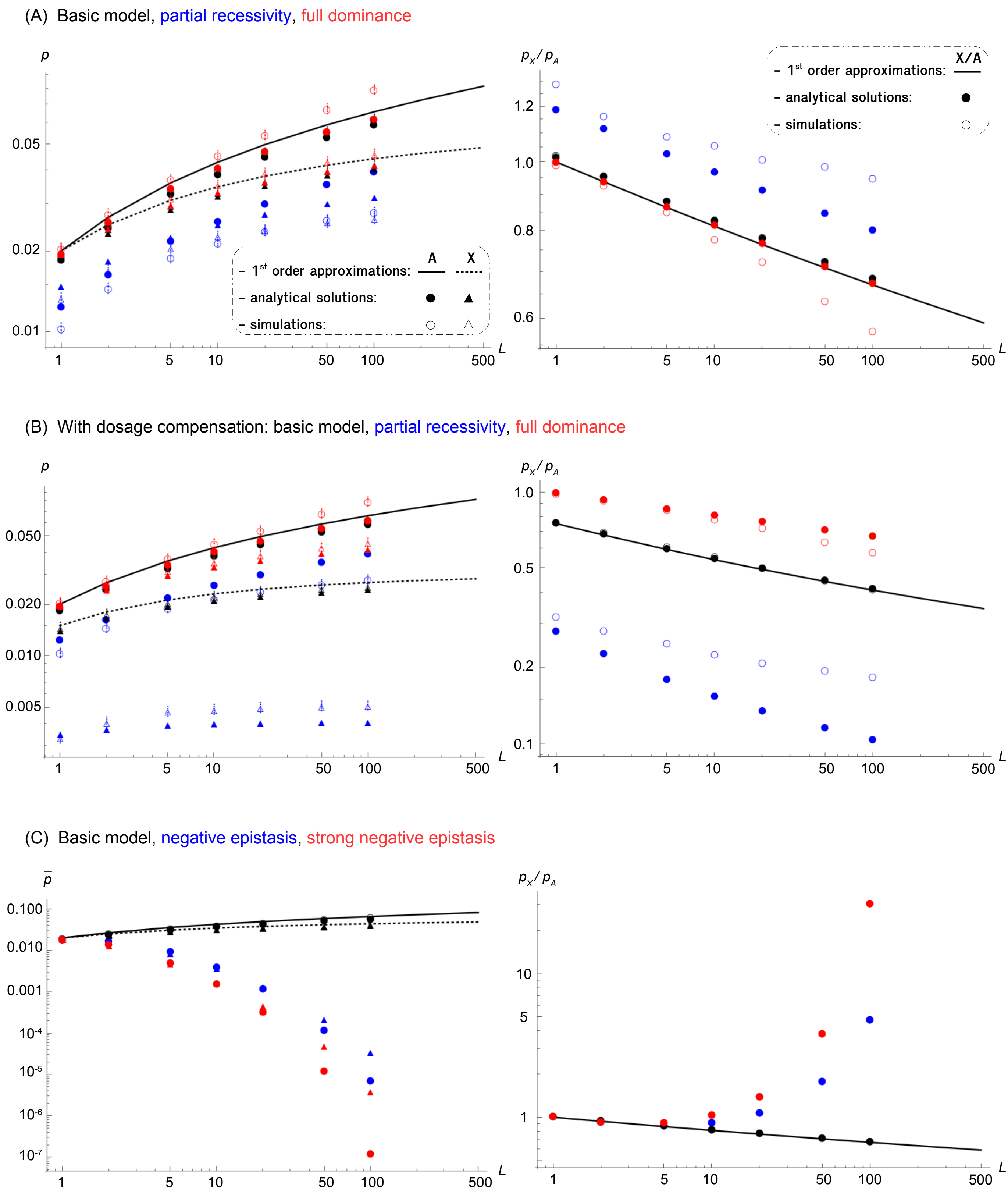


**Figure S5. Effect of model extensions on average equilibrium frequencies**

Average frequency of the deleterious alleles in the recipient species (**A**utosomal: $\overline{p}_A$; **X**-linked: $\overline{p}_X$; their ratio: $\frac{\overline{p}_X}{\overline{p}_A}$) plotted against the number of selected loci on the genomic block, $L$. **(A)** Dominance of the deleterious alleles. Results are shown for a model with co-dominance ($h = 0.5$, $s_{hom}L = 0.1$, black), and models with partial recessivity ($h = 0.1$, $s_{hom}L = 0.5$, blue) and full dominance ($h = 1.0$, $s_{hom}L = 0.05$, red). Note that eq. (4) (lines) is independent of $h$ as long as $sL, cL << 1$. **(B)** Dominance of the deleterious alleles with dosage compensation. Same as in (A) but with $s_M L = \frac{0.05}{h}$ for sex-linked alleles. Blue: $h = 0.1$, $s_{hom}L = 0.5$, $s_M L = 0.5$ for sex-linked alleles, $m_X/S_X = 0.005$, $\theta_X = 6$. Red: $h = 1.0$, $s_{hom}L = 0.05$, $s_M L = 0.05$ for sex-linked alleles, $m_X/S_X = 0.02$, $\theta_X = 1.5$. **(C)** Epistasis between selected loci. Results are shown for the basic model (i.e. multiplicative fitness: $\varepsilon = 1$, black) and models with negative epistasis (weak: $\varepsilon = 0.5$, blue; strong: $\varepsilon = 0.1$, red) in which a fraction $\beta$ of the loci acts epistatically and a fraction $1 - \beta$ acts multiplicatively. Note that we fixed $\beta = 1$, and that simulations were not computed. Parameter values for the basic model are: $h = 0.5$, $m = 0.001$, $cL = 0.05$, $sL = 0.05$, $m_X/S_X = m_A/S_A = 0.02$, $\theta_X = 1.5$, $\theta_A = 1$ and $N = 10^5$ (simulations). With dosage compensation, values are the same for the basic model, except that: $s_M L = 0.1$ for sex-linked alleles, $m_X/S_X = 0.015$, $\theta_X = 2$. Other details match Figure 1.

(A) Basic model, partial recessivity, full dominance

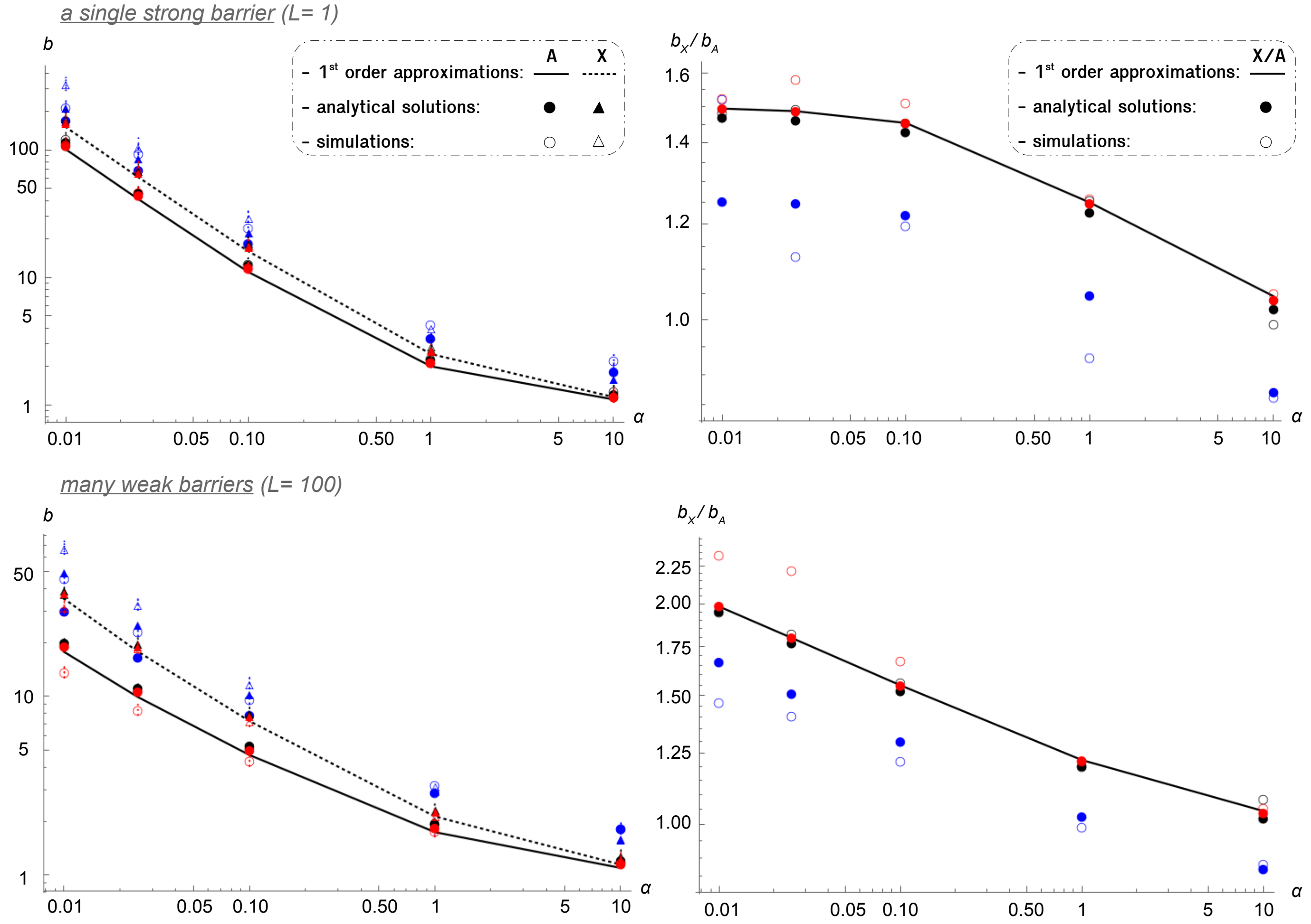


1640

1641

(B) With dosage compensation: basic model, partial recessivity, full dominance

a single strong barrier (L= 1)

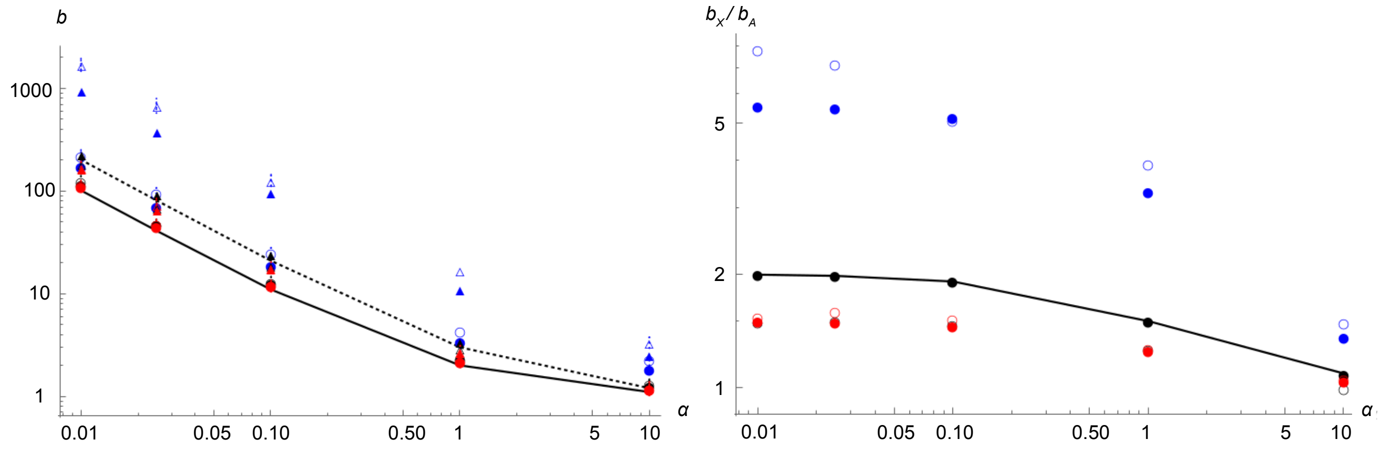


many weak barriers (L= 100)

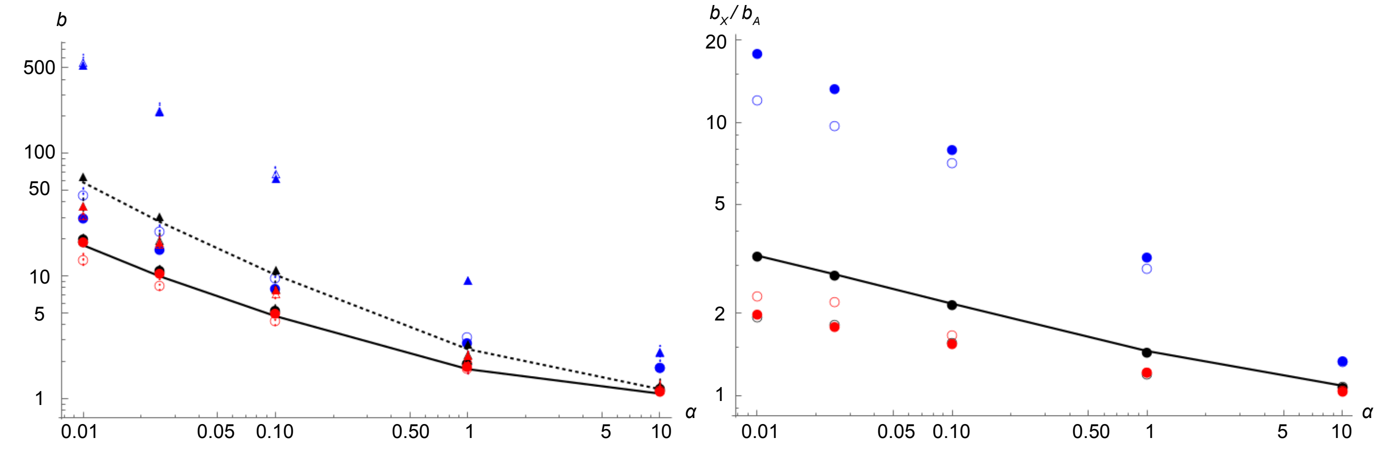


1642

1643

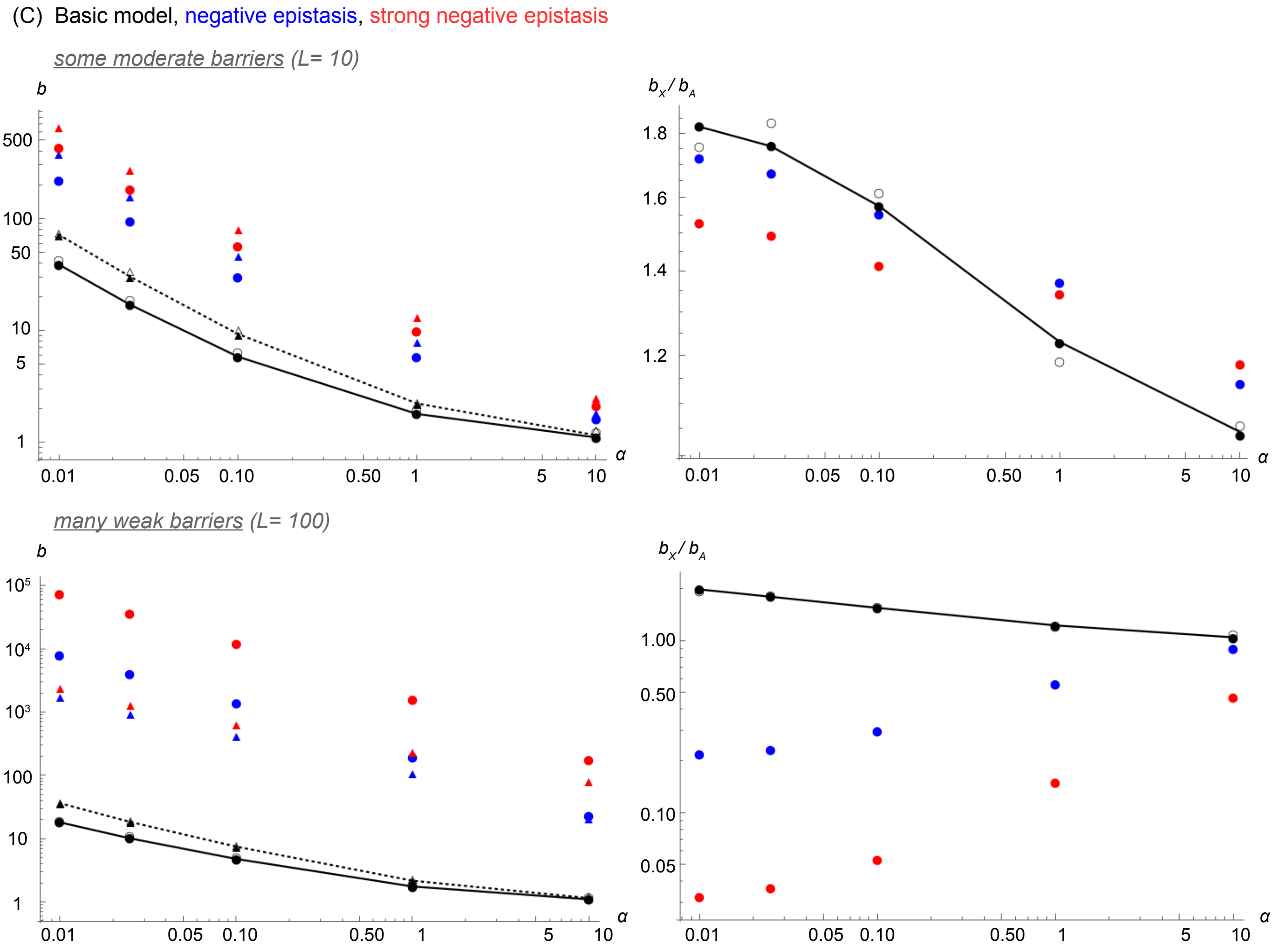


**Figure S6. Effect of model extensions on barrier strength at the neutral marker**

Barrier strength at the neutral marker (**A**utosomal: $b_A$; **X**-linked: $b_X$; their ratio: $\frac{b_X}{b_A}$) plotted against its proximity to the nearest selected locus, $\alpha$. **(A)** Dominance of the deleterious alleles ($h = 0.5$, black; $h = 0.1$, blue; $h = 1.0$, red). **(B)** Dominance of the deleterious alleles with dosage compensation. Same as in (A) but with $s_M L = \frac{0.05}{h}$ for sex-linked alleles. **(C)** Epistasis between selected loci (multiplicative fitness: $\varepsilon = 1$, black; weak negative epistasis: $\varepsilon = 0.5$, blue; strong negative epistasis: $\varepsilon = 0.1$, red). Other details match Figure 2 and Figure S5.

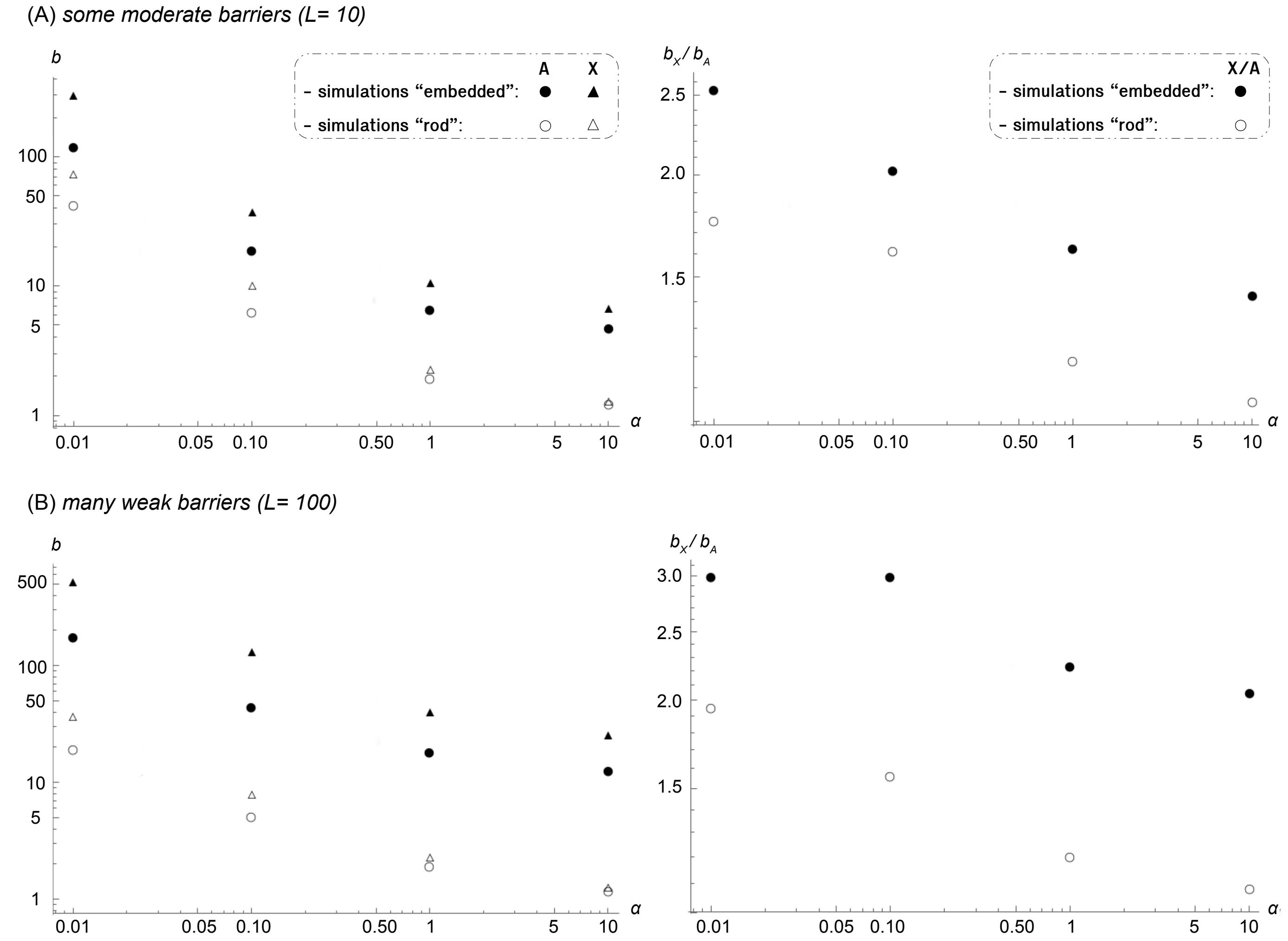


**Figure S7. Effect of the “rod” vs “embedded” configurations on barrier strength at the neutral marker**

Barrier strength at the neutral marker (**A**utosomal: $b_A$; **X**-linked: $b_X$; their ratio: $\frac{b_X}{b_A}$) plotted against its proximity to the nearest selected locus (or to each of the two surrounding loci), $\alpha$. In **(A)** the neutral marker is linked to ten moderately selected loci ($L = 10$); while in **(B)** it is linked to one hundred weakly selected loci ($L = 100$). Filled symbols show results of individual-based simulations for the case of a marker embedded in the center of the selected block, i.e. $j = k = L/2$, where $j$ selected loci lie to the left of the marker, and $k$ to the right. Empty symbols show results for a rod marker configuration, i.e. the marker is positioned at the end of the selected block ($j = L$ and $k = 0$). Parameter values are: $h = 0.5$, $m = 0.001$, $cL = 0.05$, $sL = 0.05$, $m_X/S_X = m_A/S_A = 0.02$, $\theta_X = 1.5$, $\theta_A = 1$ and $N = 10^5$ (simulations). Other details match Figure 2.

### SUPPLEMENTARY TABLES

**Table S1. Predictions for different evolutionary scenarios**

| Scenarios | | Parameters | | Predictions | |
|---|---|---|---|---|---|
| | | **autosome** | **X** | **coupling, $\theta$** | $\frac{m}{S}$ |
| Basic model | | $\beta = 0$<br>$h = 0.5$<br>$c_F = c_M = c$<br>$m_F = m_M = m$<br>$s_F = s_M = s$<br>$s_{hom,F} = s_{hom,M} = \frac{s}{h} = 2s$ | $\beta = 0$<br>$h = 0.5$<br>$c_F = c$ ; $c_M = 0$<br>$m_F = m_M = m$<br>$s_F = s_M = s$<br>$s_{hom,F} = \frac{s}{h} = 2s$ | $\theta_X = \frac{3}{2}\theta_A$ | $\frac{m_X}{S_X} = \frac{m_A}{S_A}$ |
| | for $h = 1.0$ | $s_{hom,F} = s_{hom,M} = s$ | $s_F = s_M = s_{hom,F} = s$ | as basic model | as basic model |
| | for $h = 0.1$ | $s_{hom,F} = s_{hom,M} = 10s$ | $s_{hom,F} = 10s$ | as basic model | as basic model |
| Dosage compensation | | | $s_F = s$ ; $s_M = s_{hom,F} = \frac{s}{h}$ | $\theta_X = (1 + \frac{1}{2h})\theta_A$ | $\frac{m_X}{S_X} = (\frac{3}{2+1/h})\frac{m_A}{S_A}$ |
| | for $h = 0.5$ | as basic model | $s_F = s$ ; $s_M = s_{hom,F} = 2s$ | $\theta_X = 2\theta_A$ | $\frac{m_X}{S_X} = \frac{3}{4}\frac{m_A}{S_A}$ |
| | for $h = 0.1$ | $s_{hom,F} = s_{hom,M} = 10s$ | $s_F = s$ ; $s_M = s_{hom,F} = 10s$ | $\theta_X = 6\theta_A$ | $\frac{m_X}{S_X} = \frac{1}{4}\frac{m_A}{S_A}$ |
| Achiasmy | | $c_M = 0$ | as basic model | $\theta_X = \frac{3}{4}\theta_A$ | as basic model |
| Sex-biased migration | | $m_F = km_M = m$ | $m_F = km_M = m$ | as basic model | $\frac{m_X}{S_X} = \frac{(2+4k)}{(3+3k)}\frac{m_A}{S_A}$ |
| | for $k = 3$ (♀-biased) | $m_F = 3m_M = m$ | $m_F = 3m_M = m$ | as basic model | $\frac{m_X}{S_X} = \frac{7}{6}\frac{m_A}{S_A}$ |
| | for $k = 1/3$ (♂-biased) | $m_F = \frac{m_M}{3} = m$ | $m_F = \frac{m_M}{3} = m$ | as basic model | $\frac{m_X}{S_X} = \frac{5}{6}\frac{m_A}{S_A}$ |
| Sex-biased selection | | $s_F = ks_M = s$ | $s_F = ks_M = s$ | $\theta_X = \frac{(2k+1)}{(k+1)}\theta_A$ | $\frac{m_X}{S_X} = \frac{(3+3k)}{(2+4k)}\frac{m_A}{S_A}$ |
| | for $k = 2$ (♀-biased) | $s_F = 2s_M = s$ | $s_F = 2s_M = s$ | $\theta_X = \frac{5}{3}\theta_A$ | $\frac{m_X}{S_X} = \frac{9}{10}\frac{m_A}{S_A}$ |
| | for $k = 1/2$ (♂-biased) | $s_F = \frac{s_M}{2} = s$ | $s_F = \frac{s_M}{2} = s$ | $\theta_X = \frac{4}{3}\theta_A$ | $\frac{m_X}{S_X} = \frac{9}{8}\frac{m_A}{S_A}$ |

The predicted values of $\theta$ and $\frac{m}{S}$ are derived from the analytical solution in eq. (4) (which assumes $sL$ and $cL \ll 1$), and so they are independent of the dominance coefficient ($h$). See Table 1 for the significance of model parameters.
